# Non-coding function for mRNAs in Focal Adhesion Architecture and Mechanotransduction

**DOI:** 10.1101/2021.10.04.463097

**Authors:** Liana Boraas, Mengwei Hu, Lauren Thornton, Charles E. Vejnar, Gang Zhen, Antonio J. Giraldez, Christine Mayr, Siyuan Wang, Stefania Nicoli

**Author notes:** These authors contributed equally (Liana Boraas, Mengwei Hu).

## Abstract

Messenger RNA (mRNA) compartmentalization within the cytosol is well-recognized as a key mechanism of local translation-mediated regulation of protein levels, but whether such localization could be a means of exercising non-coding mRNA function is unknown. Here, we explore non-coding functions for mRNAs associated with focal adhesions (FAs), cellular structures responsible for mediating cell adhesion and response to changes in the extracellular matrix (ECM). Using high-throughput single molecule imaging and genomic profiling approaches, we find that mRNAs with distinct sequence characteristics localize to FAs in different human cell types. Notably, ∼85% of FA-mRNAs are not translationally active at steady state or under conditions of FA dissolution or activation. Untranslated mRNA sequences are anchored to FA based on their functional states by the RNA binding protein, G3BP1, forming biomolecular granules. Removing RNA or G3BP1, but not blocking new polypeptide synthesis, dramatically changes FA protein composition and organization, resulting in loss of cell contractility and cellular ability to adapt to changing ECM. We have therefor uncovered a novel, non-coding role for mRNAs as scaffolds to maintain FA structure and function, broadening our understating of noncanonical mRNA functions.

## Main Text

Mounting evidence supports the notion that intracellular localization and organization of mRNAs is a prevalent mechanism to regulate protein production and function in numerous cellular processes^1^ and that the dysregulation of such control results in multisystemic diseases^2,3^. Initially discovered in neurons and highly polarized cells such as oocytes^4^ or migrating embryonic fibroblasts^5^, RNA localization is now recognized as a wide-spread phenomenon in many cell types.

One of the main mechanisms to localize RNAs is the formation of hydrogel or phase-separated droplets where RNA Binding Protein (RBPs) form complexes with one or multiple RNA molecules, forming ribonucleoprotein (RNP) granules^6^. Cytosolic mRNAs can also be enriched within several cellular compartments^7^, such as the endoplasmic reticulum or the mitochondria, to form messenger RNPs and mediate local translation that facilitates key protein-protein interactions and/or signaling^8-10^. In somatic cells, messenger RNP granules can also function to keep mRNA translationally inactive (processing-p-bodies) or to sequester it after translation is temporarily arrested by stress conditions (stress granules)^11,12^. Nevertheless, in homeostasis most cytosolic and compartmentalized mRNA are recognized to be translated and their control serves to locally regulate protein levels.

Even though localized mRNAs have proven to be a critical mechanism to sustain cellular homeostasis, fundamental understanding of the spectrum of biological functions underlying these mRNA foci is lacking. Filling this knowledge gap will establish a comprehensive framework of how mRNAs regulate cell functions beyond their canonical role in encoding proteins.

Here we took an unbiased approach to identify the nature and function of mRNA localized at focal adhesions (FAs). FAs are multifunctional organelles that connect via cell surface receptors, such as integrins, the actin cytoskeleton to the extracellular matrix (ECM). FAs respond continuously to mechanical cues, such as matrix protein composition and stiffness^13,14^, and are essential for cellular processes that, if lost, can contribute to numerous pathologies including cancer, defects in wound healing and fibrotic diseases^15^. To meditate mechanoresponses, FAs require an arsenal of ∼200 proteins, whose composition and organization change from smaller, circular nascent structures to mature adhesive complexes 2 μm wide and 3-10 μm long in size ^16-18^. FAs number, morphology and cellular distribution reflect the contractile state of the cell, which is key to transduce cell-matrix interactions^19,20^. Several RBPs have being identified as binding partners of FA proteins^21-26^, and mRNAs have been found localized and translated in multiple types of cell protrusions and lamellipodia, which are notably enriched in FAs^5,27-31^. However, potential non-canonical functions of mRNAs at FAs have not been addressed.

Here, we employed a hydromechanical method to isolate intact FAs from the rest of the cell body (Fig.1A and B and fig. S1A)^32^. Total RNAs were collected from whole cells and FAs isolates from human dermal fibroblasts (HDFs) and human umbilical vein endothelial cells (HUVECs), two different adherent cell type^33,34^, and polyadenylated mRNA was profiled using RNA-sequencing (fig. S1B). We identified 236 and 119 mRNAs that were significantly enriched in HUVEC and HDF FAs, respectively, and that 57 were shared between the two cell types (Fig. 1C and 1D and Table S1). Interestingly, we found that ∼80% of the shared FA mRNAs encoded for proteins important for cellular processes controlled by FAs, such as proteins involved in actin, cytoskeleton and cell adhesion protein binding. In addition, the FA mRNAs systematically exhibited higher overall and 3’UTR sequence length, number of AU-rich elements, and intermolecular RNA-RNA interactions, as compared to the whole cell mRNAs control^35^ (Fig. 1D and E, fig. S1C and D). Therefore, mRNAs localized at FAs are conserved among human cells and possess distinctive sequence characteristics.

**Figure. 1.**
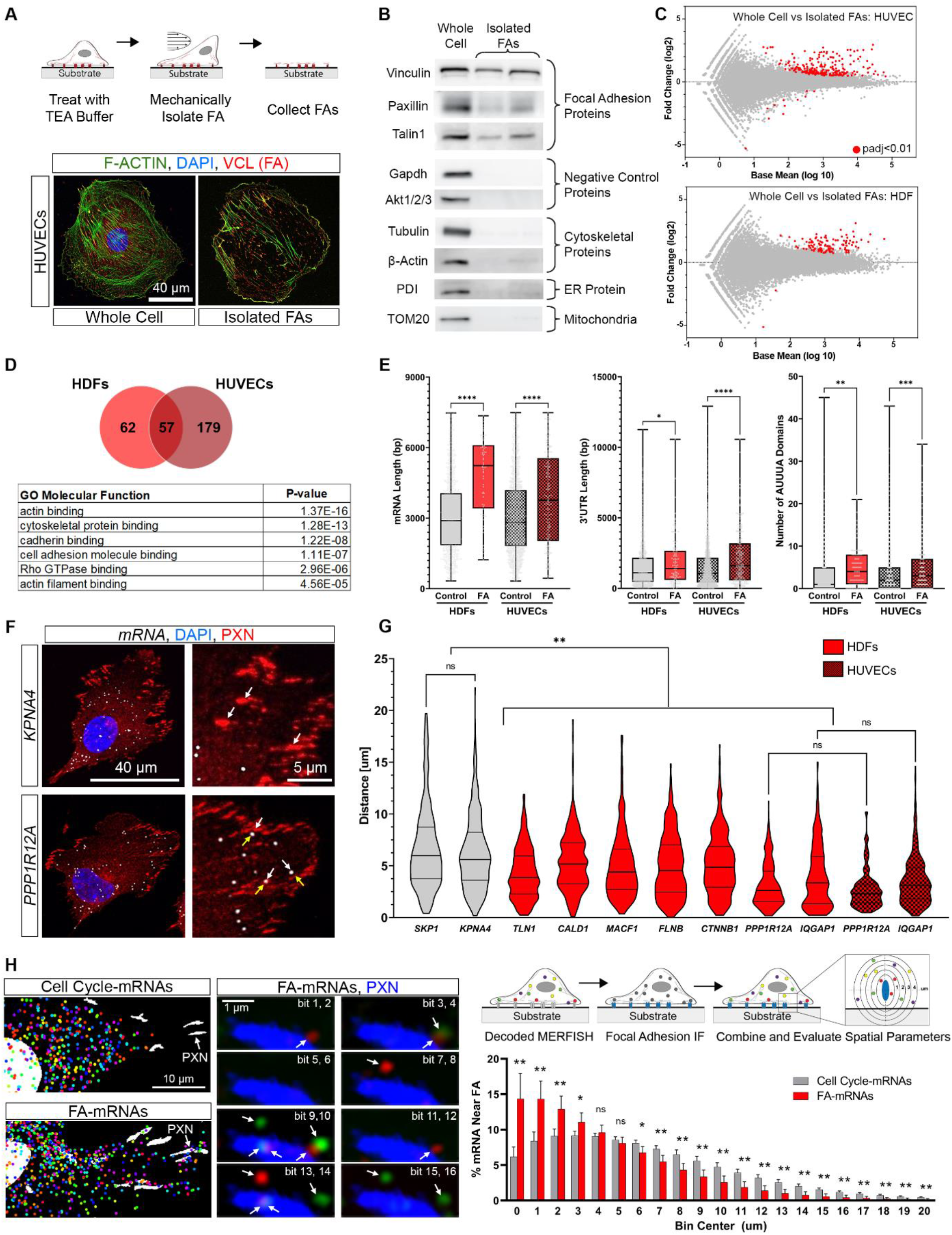
A distinct subpopulation of mRNAs localized to focal adhesions. (**A**) (Top) Schematic of focal adhesion (FA) isolation protocol. (Bottom) Representative immunofluorescence images of HUVECs after FA isolation. Cells were stained for F-ACTIN (phalloidin), Vinculin (VCL), and DAPI. (**B**) Western blots of whole cell or isolated FA samples for FA proteins (Vinculin, Paxillin, Talin1), negative control proteins not found in FA (Gapdh, Akt1/2/3), cytoskeletal proteins (Tubulin, B-actin), ER protein (PDI) or a mitochondria protein (TOM20). (**C**) mRNA sequencing results for whole cell vs isolated FAs in HUVECs and HDFs. Data plotted as average expression vs Log2 fold-change with a padj<0.01 for significantly localized mRNAs with four independent replicates for each group. (**D**) (Top) Venn diagram for mRNAs significantly localized to FAs in HDFs and HUVECs. (Bottom) Top GO Molecular Function terms for shared mRNAs between the two cell types. (**E**) Box plots of overall mRNA length, 3’UTR length, and number of AUUUA sequences in the 3’UTR for control mRNAs (grey) and FA localized mRNA (red) in HDFs (solid) or HUVECs (checkers). (**F**) Representative smFISH + immunofluorescence images of HDFs where cells were probed for mRNA and stained for Paxillin (PXN, white arrows) and DAPI. Yellow arrows indicate regions of mRNA near FA. (**G**) Quantification of distance from an mRNA to its closest FA for control (grey) and FA localized mRNA (red) in HDFs (solid) or HUVECs (checkers). Each violin plot represents quantification from 10-12 cells with a total of 156-484 mRNA analyzed per mRNA species (**H**) (Left) Composite MERFISH images for Cell Cycle and FA mRNAs after all hybridization rounds with FAs marked by paxillin (PXN, white arrows) in HDFs. Raw fluorescent images from eight rounds of hybridization where white arrows mark a positive signal for a single molecule spot and can be identified using the assigned binary code under perfect match criteria. (Right, Top) Schematic of method to evaluate localization of mRNA near FAs. (Right, Bottom) Histogram of the percent of all mRNA localized within a given distance to the FA edge for Cell Cycle-mRNAs (grey) or FA-mRNAs (red) and the two distributions are statistically significant from one another (****P < 0.0001). Each bin is the average percent near FA for every mRNA in each category across 5 independent replicates with a total of 239 FOV and a total of 1,932 cells per group. For all graphs, significance is represented as not significant (n.s.) P > 0.05, *P ≤ 0.05, **P ≤ 0.01,***P ≤ 0.001, ****P ≤ 0.0001.

Using single molecule fluorescence *in situ* hybridization (smFISH), we verified that FA-localized mRNAs (FA-mRNAs) were found near the center of Paxillin (PXN) proteins, a core FA component, of both HDFs and HUVECs, in comparison to mRNAs identified in the whole cell (Fig. 1F and G, fig. S1E and Table S2). To further spatially resolve FA-mRNA localization with high accuracy, we used the multiplexed error-robust FISH (MERFISH)^36^ approach. We designed a library of oligonucleotide probes to visualize mRNAs commonly present in HDF and HUVEC FAs and a control library of mRNAs encoding for cell cycle proteins, unrelated to our identified FAs mRNAs (Table S2). To visualize mRNAs localization to FAs, we performed MERFISH using PXN as co-staining (Fig. 1H). mRNA localization over five reproducible experiments found that the distribution of FA-mRNAs was significantly closer to FAs in comparison to cell cycle mRNAs and that this enrichment was independent of the mRNAs’ overall abundance within cells (p<0.0001, Fig. 1H and fig. S1F, G, and H). In all, these data suggest that there is a cohort of mRNAs specifically localized to FAs.

Since mRNAs at FAs encoded for proteins relevant to FA function, we tested whether translation may be the main functional reason for their localization. HDF and HUVEC FA isolates were prepared for ribosome profiling and proteomics analysis to define both mRNA translation and protein localization, respectively (Fig. 2A). Notably, in HDFs, 83% of FA-associated mRNAs were not detected as proteins in the FA, nor were detected as being actively translated, ribosome-associated transcripts. 87% of FA-associated mRNAs in HUVECs exhibited the same lack of protein product and association with active translation (Fig. 2B and Table S3). To confirm these observations, the nascent polypeptide of putative “translated” or “untranslated” FA-mRNAs was detected via puromycylation coupled with proximity ligation (Puro-PLA)^37^. We first tested mRNA translation by sampling “translated” mRNA species in HDF FAs that were identified together with their proteins and ribosome protected fragments (∼27%, Table S3). We found that both mRNA and the forming polypeptide localized at FA, confirming their translational status (Fig. 2C, 2D and fig. S2A). Next, we tested FA mRNAs categorized as “untranslated” (∼83%, Table S3). While we detected mRNAs localized at FAs for all the tested mRNA species, we found 35 to 90% less mRNA-newly polypeptide FA localization, supporting that most FA-mRNAs were not translationally active (Fig. 2C and 2D).

**Figure 2.**
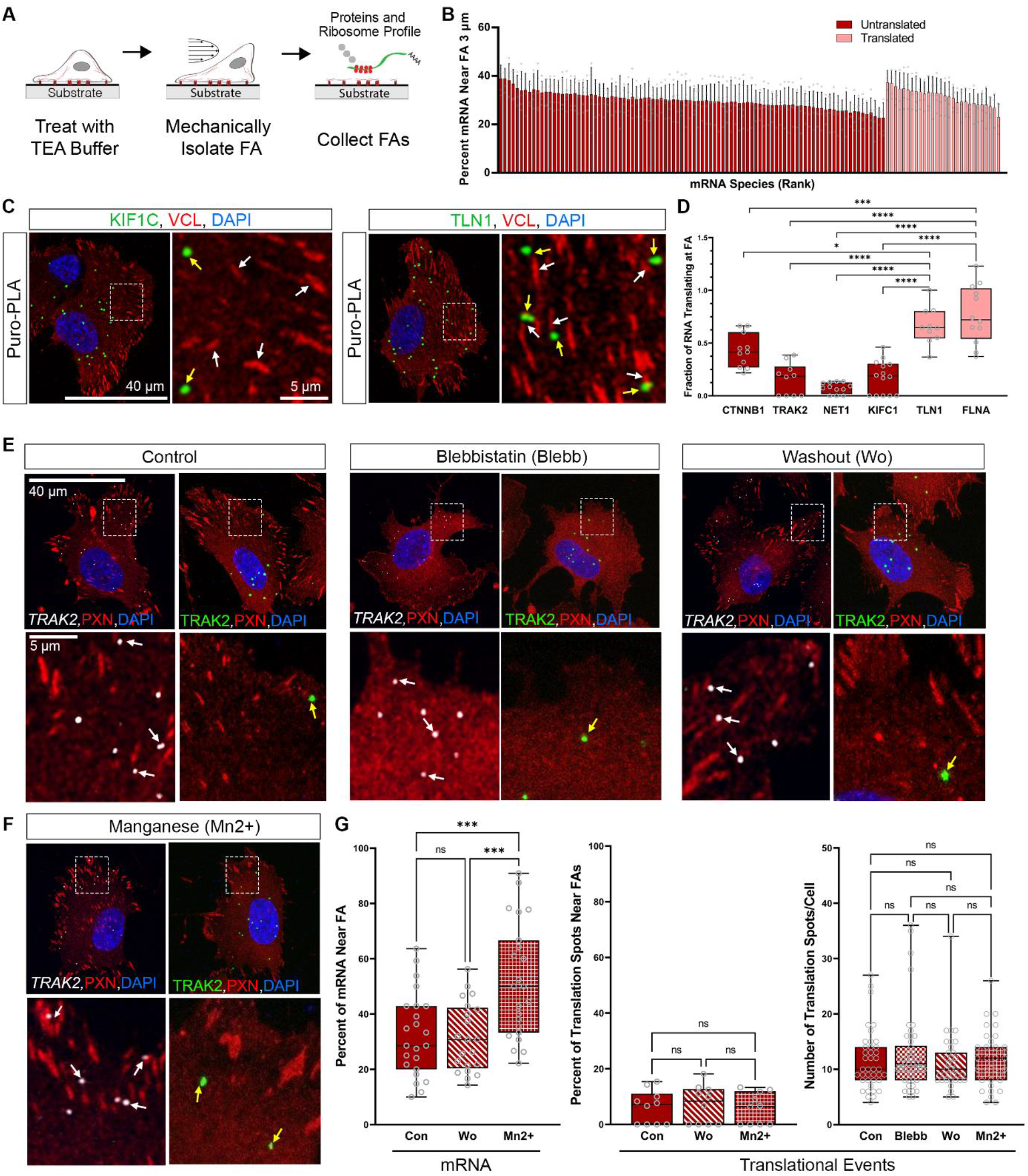
The majority of FA mRNAs are translationally inactive. (**A**) Schematic of focal adhesion (FA) isolation for proteomics and ribosome profiling. (**B**) Untranslated (dark red) or translated (light red) mRNAs ranked by the percent mRNA within 3 μm of FAs from MERFISH quantification in Fig. 1H. (**C**) Representative Puro-PLA images for KIF1C (untranslated) and TLN1 (translated) with spots of translation (green, yellow arrows), vinculin (VCL, red, white arrows) and DAPI (blue) shown in HDFs. (**D**) Fraction of RNA translating near FA (defined at <2.2 μm from FA center) for untranslated (dark red) or translated (light red) mRNAs (n=10-15 cells per mRNA species with a total of ∼10-100 spots per cell depending on the species). (**E**) Representative images for control, blebbistatin treated (25 uM for 30 minutes), and washout cells (30 minutes after removal of blebbistatin), (**F**) as well as manganese treated cells (3 uM for 20 minutes). All cells were stained for focal adhesions (Paxillin, PXN) and probed for either mRNA (smFISH, white arrows) or translational events (Puro-PLA, yellow arrows). Doted boxes depicted on top panels represent the region highly magnified in the bottom panels. (**G**) Box plots for the percent of mRNA near FAs (n=20-25 cells), percent translational events near FAs (n=10-15 cells) (both defined at <2.2 μm from FA center), and the overall number of translational events per cell (n=45 cells). For all graphs, significance is represented as not significant (n.s.) P > 0.05, *P ≤ 0.05, **P ≤ 0.01,***P ≤ 0.001, ****P ≤ 0.0001.

We next tested whether the translation of FA-mRNAs was dependent on FA function. We thus performed smFISH and puro-PLA in HDFs under conditions that induce de-novo FA formation or activate pre-existing FAs. First, cells were treated for 30 min with a myosin II inhibitor, Blebbistatin (Blebb) to disassemble FAs, before the inhibitor was washed out to rapidly reinstate FA formation^38^. Under these conditions, we found that *TRAK2*, an untranslated FA-mRNA, did not form its polypeptide at new FA sites nor in other cellular compartments when compared to untreated cells (Fig. 2E and 2G). Second, we stimulated pre-existing FAs in HDFs via manganese (Mn2+)-induced integrin maturation^39^. Notably, we detected an increased number of *TRAK2* mRNAs near larger and Mn2+-activated FAs but failed to detect a correspondent increase in puro-PLA staining at FAs or within the whole cells (Fig. 2F and 2G). Other untranslated FA-mRNAs behaved similarly to *TRAK2*, whereas putative translated FA-mRNAs showed changes in protein translation depending on FA dynamics (fig. S2B and C). All together, these data suggest that much of the mRNA localized at FAs is untranslated in both steady state and activated FAs.

In order to continually interpret cell-matrix interactions, FAs exist in different dynamics states, characterized by diverse morphologies, cellular distribution, and number^19,20^. Here, we sought to understand if mRNAs distribution is related to the functional states of FAs. We used PHATE, a method that provides a representation of a high-dimensional dataset^40^, to visualize the states of FAs defined by their nearby mRNA sequences. Strikingly, we found that untranslated mRNAs were differentially localized with four main subtypes of Paxillin (PXN)-positive FAs in HDF (FA-clusters), whereas translated FA-mRNAs were not (Fig. 3A and fig. S3A). Of the fours FA-clusters, FA-cluster 3 and 0 have the most distinct characteristic from each other, where FA-cluster 3 were significantly smaller, more circular, and further away from the nuclei than FA-cluster 0 (Fig. 3B). To confirm these observations, we selected mRNAs markers (Fig. 3C) of these two distinct FA-clusters and performed smFISH analysis. We found *KIF1C* and *TRAK2*, which were predicted by PHATE to localize near FA-cluster 3, were indeed detected at FAs with smaller size, higher circularity, and longer distance from the nucleus (Fig. 3D and fig. S3B). In contrast, *STEAP3* and *ZMPSTE24* were predominantly localized to larger FAs under the nucleus corresponding to FA-cluster 0 (Fig. 3D and fig. S3B). Mn2+-integrin-dependent activation of FA-cluster 3 and 0 resulted in an increased percentage of the respective mRNA sequence markers (Fig. 3E). Overall untranslated mRNAs localization is tightly associated with key functional FA states.

**Figure 3.**
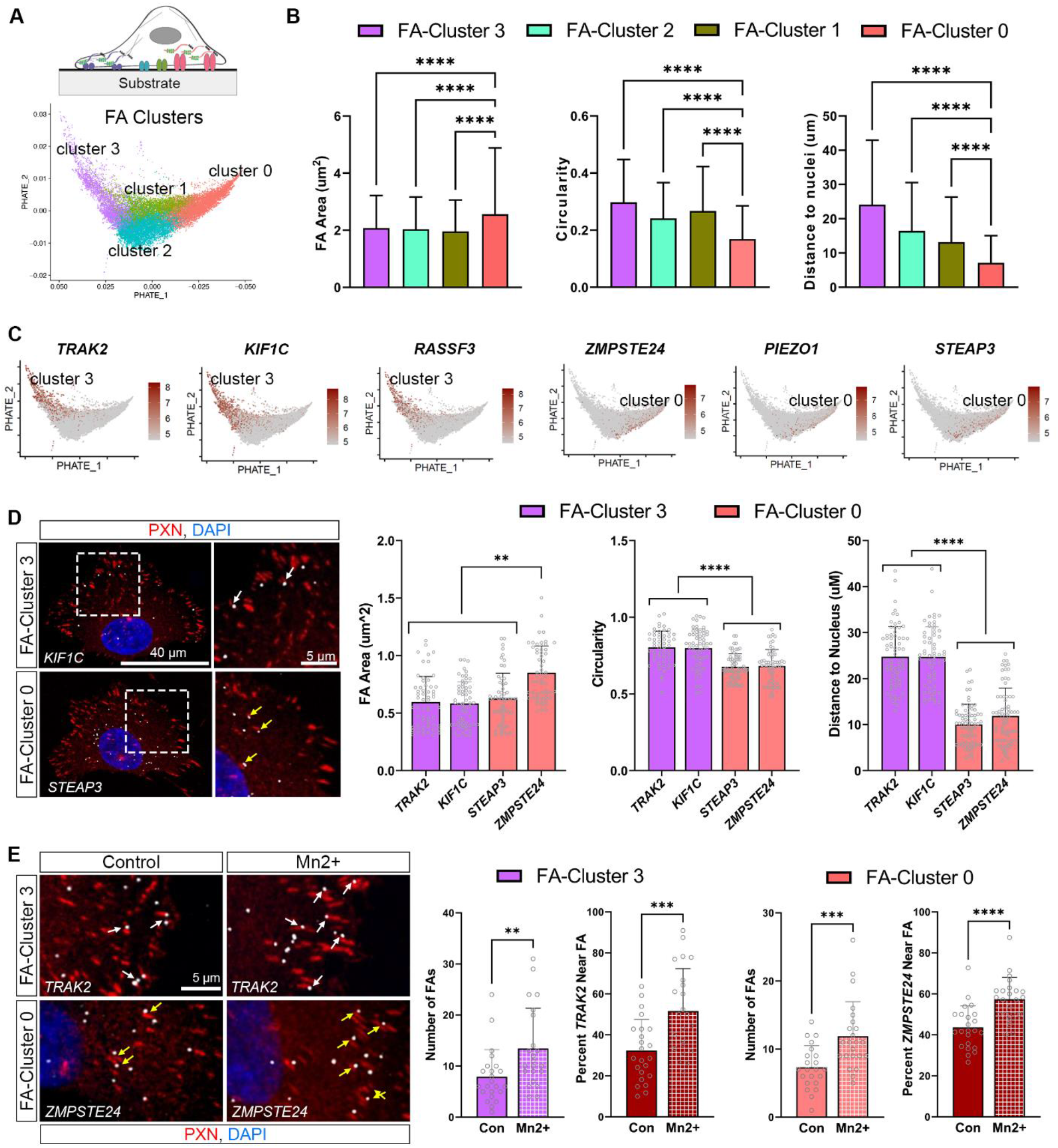
Specific untranslated mRNAs associate with distinct FA states. (**A**) Schematics (top) and visualization of PHATE analysis of FAs based on their associated (within 2.2 μm to FAs) mRNAs visualized via MERFISH (bottom). (**B**) Properties of FAs within each cluster (FA size, circularity, and distance to nucleus) from MERFISH analysis of the mRNAs are shown as mean ± STDEV. (**C**) Feature plots of defining markers for cluster 0 (*ZMPSTE24, STEAP3, PIEZO1*), cluster 3 (*TRAK2, KIF1C, RASSF3*). (**D**) (Left) Representative images of cluster 3 or cluster 0 mRNA (white), FA (PXN, red), and DAPI. White arrows indicate FAs in cluster 3 and yellow arrows indicate FAs in cluster 0. Doted boxes depicted on left panels represent the region highly magnified in the right panels. (Right) Mean ± STDEV quantification of size, circularity, and distance to nucleus for FAs near (defined at <2.2 μm from FA center) the given mRNA marker (n=60-75 FA per group). (**E**) Representative images of mRNAs (cluster 3, *TRAK2*; cluster 0, *ZMPSTE24*) and FAs (PXN, red) in control and manganese treated cells (3 μM for 20 minutes). Quantification of the number of FA per cluster or the percent mRNA near FAs (defined at <2.2 μm from FA center) are show as mean ± STDEV (n=20-25 cells). Cells were treated with either sham solvent or 3 uM Mn2+ for 20 minutes. For all graphs, significance is represented as not significant (n.s.) P > 0.05, *P ≤ 0.05, **P ≤ 0.01,***P ≤ 0.001, ****P ≤ 0.0001.

Since our data suggest that mRNA sequences are associated to FA function rather that the proteins they encode, we set out to investigate this unexpected non-coding role for mRNA at FAs. We first assayed FA states in HDFs treated with ribonuclease (RNase A) for 10 min to remove RNA, as compared to cells treated with Cycloheximide (CHX) to block mRNA-protein synthesis (Fig. 4A and fig. S4A and B). We found RNase A-treated cells exhibited a drastic diminishment of FA size, an increase in circularity and a slightly decrease in FA number, as compared to untreated or CHX-treated cells (Fig. 4B and 4C). Furthermore, proteomics analysis of FA isolates from control, RNase A- and CHX-treated cells supported the observation that FAs protein composition was most significatively changed in cells after RNA depletion (Fig. 4D and Table S4). Thus, mRNAs have a distinct function from translation in the regulation of FAs protein composition and consequently morphology.

**Figure 4.**
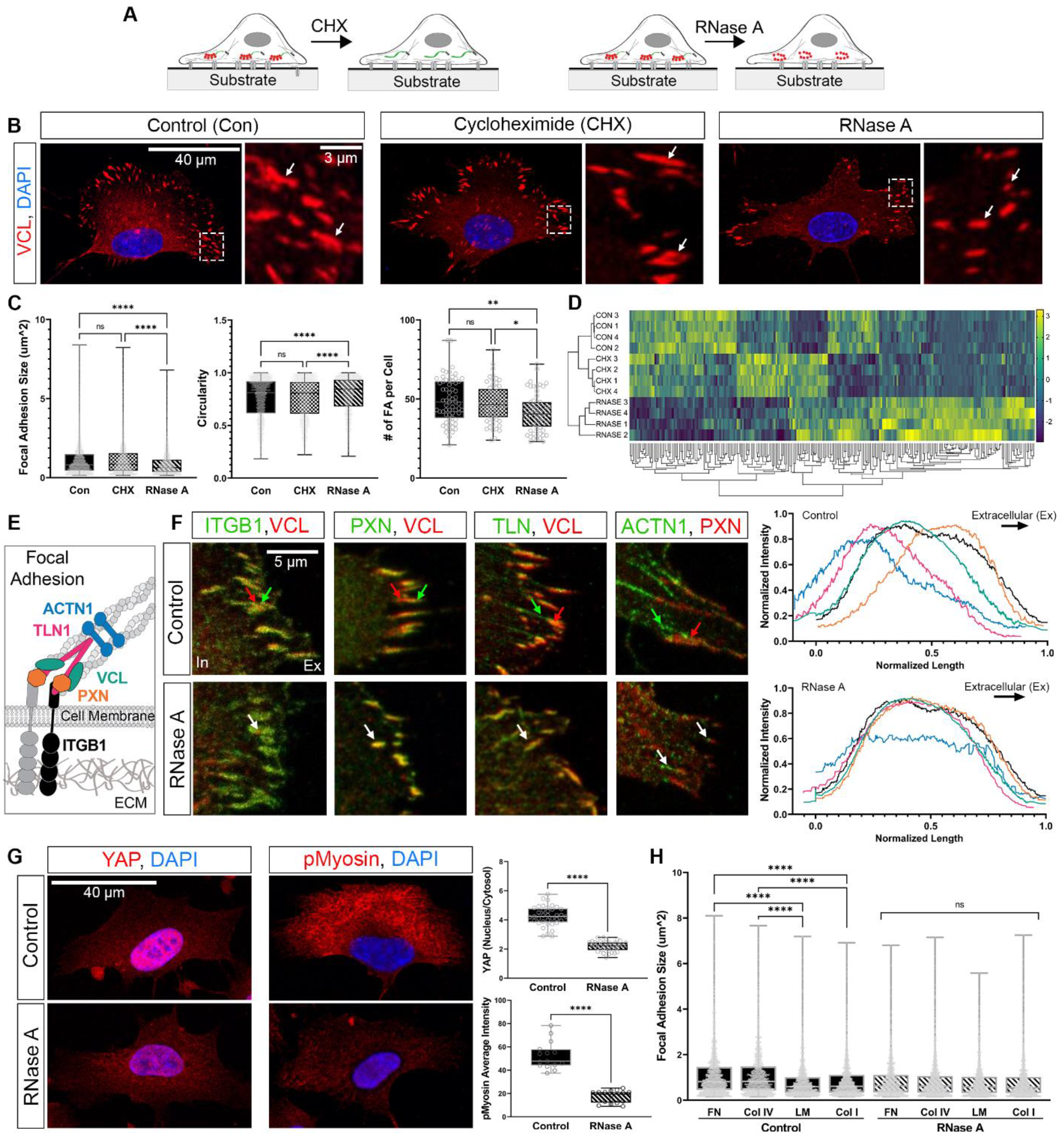
Removal of RNA, but not translation, alters FA protein organization and function. (**A**) Schematic of cyclohexamide (CHX, 100 μg/mL for 10 minutes) and RNase A treatment (1 mg/ml for 10 minutes) on cultured cells. (**B**) Representative images of FAs marked by vinculin (VCL, red, white arrows) in control, CHX, or RNase A treated HDFs. Doted boxes depicted on left panels represent the region highly magnified in the right panels. (**C**) Box plots of FA size, circularity, and overall number of FA per cell in control and treated cells. For each group tested, 58-63 cells were quantified with ∼50 FA per cell. (**D**) Heat map of differentially expressed proteins from liquid chromatography–mass spectrometry analysis of isolated FAs in control and treated cells (color legend represent z-scores, clustering based on z-score, n=4 independent replicates per group). (**E**) Schematic of FA protein organization with core proteins ITGB1 (black), PXN (orange), VCL (green), TLN1 (pink), and ACTN1 (blue) shown. (**F**) Representative images of FA proteins in control and RNase A treated cells with the intracellular space (In) and extracellular space (ex) indicated. Red and green arrows in control indicate the respective protein organization, while white arrows in the bottom panel indicate disruption of this organization. Protein intensity was quantified for each core FA protein. Length of each FA was normalized using VCL or PXN and was also used as a reference to align the other core proteins. Expression intensity was normalized to max intensity. 25 FAs per pair of proteins was analyzed from at least 5 different cells for a total of 100 FAs per treatment group. (**G**) Representative images of YAP and pMyosin expression in control and RNase A treated cells (red) and quantification of YAP nuclear to cytoplasmic ratio or average pMyosin intensity per cell in each group (n=∼40 cells for YAP and n=∼15 cells for pMyosin quantification). (**H**) Box plot of FA size in either control or RNase A treated cells cultured on fibronectin (FN), collagen IV (Col IV), laminin (LM), or collagen I (Col I). For each group tested, ∼25 cells were quantified with ∼50 FA per cell. For all graphs, significance is represented as not significant (n.s.) P > 0.05, *P ≤ 0.05, **P ≤ 0.01,***P ≤ 0.001, ****P ≤ 0.0001.

To further investigate this point, we analysed the effect of depleting RNAs on FA protein architecture. FAs are vertically organized into strata, composed of a basal layer of integrin cytoplasmic tails and Paxillin, a middle layer of Talin and Vinculin and an upper-most Actin-regulatory layer that is essential to connect the matrix to the cytoskeleton^41^ (Fig. 4E). Remarkably, the abundance of these core proteins was not de-regulated in the FAs of RNase A-treated HDFs, but the protein organization of the middle and the actin-regulatory layer of the FA strata was completely lost (Fig. 4F and fig. S4C). Furthermore, RNase A-treated cells showed a significant decrease in the nuclear localization of the mechanosensitive transcription factor Yes-associated protein (YAP)^42^ and in the overall level of phosphorylated myosin^43^ (Fig. 4G), both of which are consistent with the disruption of the FA protein strata and the resulting smaller and circular FAs morphology.

So far, our data suggest that mRNAs might have a non-coding role in the organization of FA protein composition and architecture, which in turns impacts FA mechanoresponses such as cell contractility. If this is the case, RNase A-treated cells would not be able to adapt to substrates consisting of different structural and compositional ECM proteins^44^. Indeed, we found that while control HDFs successfully adapted their FA size depending on the ECM protein they were cultured on (fibronectin, collagen IV, laminin, or collagen I), we found no difference between FA size in RNase A-treated cells seeded on any of the four ECM proteins tested (Fig. 4H and fig. S4D). Together, these data suggest that the mRNAs, rather than the protein they encode, are required to regulate FA proteins architecture and composition to ensure cytoskeletal contractility and ECM adaptation.

We next searched for RBPs that could explain the localization and function of our identified non-coding FA-mRNAs. We selected RBPs enriched in our proteomics analysis of HDF FA isolates vs whole cells, and depleted after treatment with RNase A, but not with CHX treatment or in control untreated cells (Fig. 4D and Table S4). Surprisingly, the top four RBPs we identified, G3BP1, G3BP2, DDX3X, and RBM3, are all components of the canonical stress granule proteome^45^, with G3BP1 being the most abundant (Fig. 5A). G3BP1 (GTPase-activating protein-binding protein 1) is a protein with intrinsically disordered regions that triggers untranslated mRNAs-dependent liquid-liquid phase separation under stress-induced translational repression^46,47^. G3BP1 or 2 function is mainly described in the context of a variety of cellular stressors such as chemotherapeutic drugs, UV irradiation, heat shock or oxidative stress; thus, its role is less recognized in isolated cells in otherwise healthy conditions. Interestingly, G3BP1 has been co-immunoprecipitated or localized with several core FA proteins in other studies, suggesting that G3BP1 could be the main protein linking the organizational function of mRNAs to the FA protein complex (fig. S5A). To test this hypothesis, we examined the localization of G3BP1 in HDFs and co-stained for PXN as a marker of FAs. We found that G3BP1 localized at FAs, forming branched and elongated granules of ∼150 nm in average diameter along the outline of PXN+ FAs (Fig. 5B, 5C and fig. S5B). G3BP2 and DDX3X proteins showed similar FA-co-localization patterns (fig. S5C). HDFs treated with RNase A for 10 minutes lost FA G3BP1+ granules relative to control untreated cells, suggesting that mRNAs are necessary for FA-co-localization (Fig. 5C).

**Figure 5.**
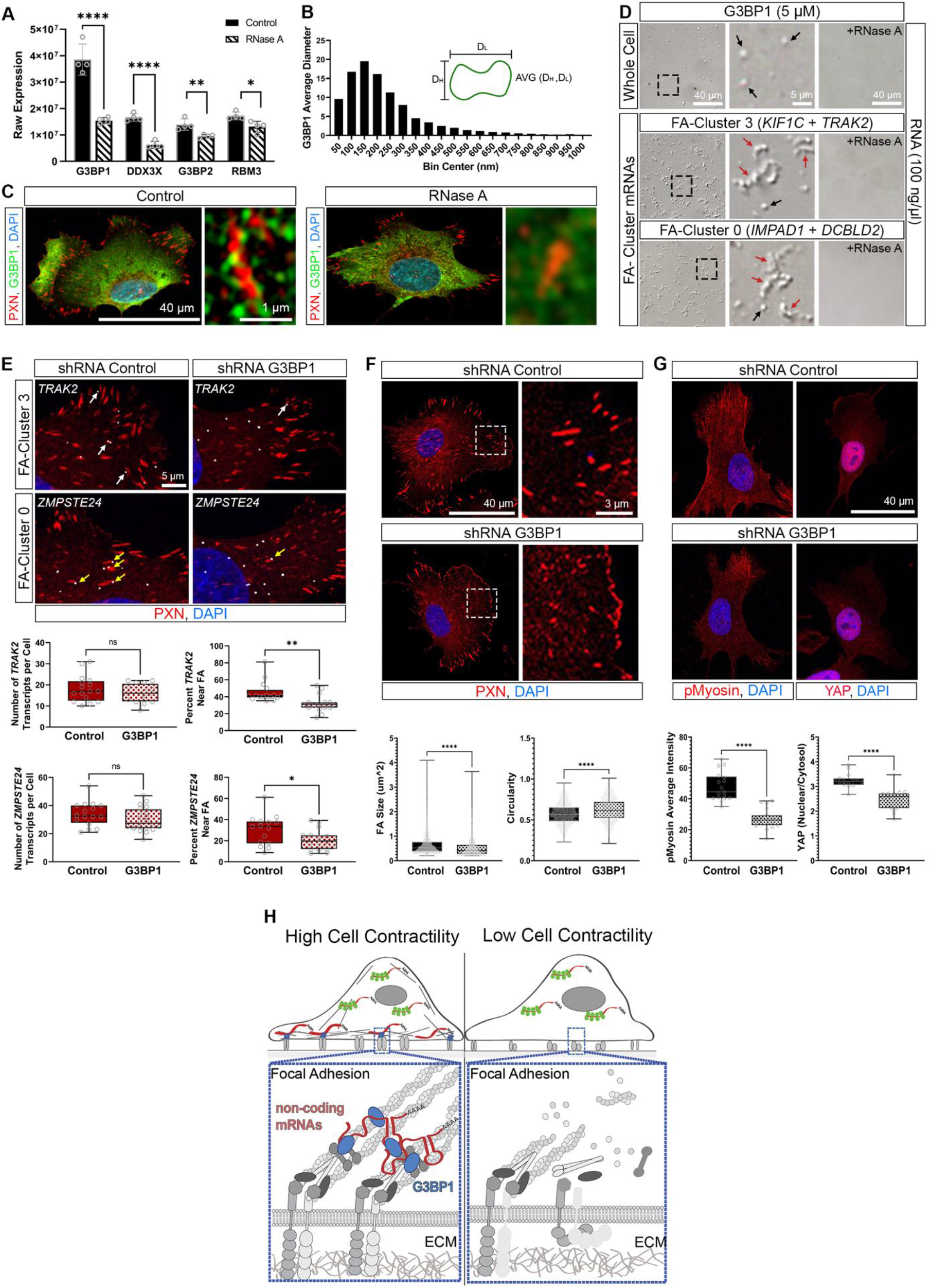
mRNA-G3BP1 granules form at FA to regulate cell contractility. (**A**) Expression of RNA binding proteins from proteomics analysis of FA isolates from control or RNase A treated HDFs (n=4). (**B**) Quantification of G3BP1 particles in control cells. ImageJ particle analysis was used to take the maximum and minimum Feret diameter and the average of the two was calculated (n=3616 clusters, data plotted as histogram, bin size = 50 nm). (**C**) Representative image of G3BP1 (green) expression in control or RNase A treated (1 mg/ml for 10 minutes) HDFs counterstained for PXN (Red) and DAPI (Blue). (**D**) DIC images of G3BP1 protein mixed with either whole cell (total) RNA or *in vitro* transcribed mRNAs with or without RNase A (1 mg/ml for 10 minutes). Black arrows indicate spherical granules and red arrows indicate branched and elongated granules. For cluster 3 *TRAK2* and *KIF1C* RNA was mixed and for cluster 0 *IMPAD1* and *DCBLD2* RNA was mixed for a final concentration of 100 ng/μl. (**E**) (Top) Representative images of mRNAs (cluster 3, *TRAK2*, white arrows; cluster 0, *ZMPSTE24*, yellow arrows) and FAs (PXN, red) in cells treated with shRNAs against G3BP1 or a non-specific control. (Bottom) Box plots of the number of mRNA transcripts per cell or the percent mRNA near FA (defined at <2.2 μm from FA center) for each cluster marker. (**F**) Representative images of FAs (PXN, red) in cells treated with shRNAs against G3BP1 or a non-specific control. Doted boxes depicted on left panels represent the region highly magnified in the right panels. Quantification of FA size and circularity shown as box plots (n=∼1500 FAs, from ∼15 cells). (**G**) Representative images of YAP and pMyosin expression in control and G3BP1 shRNA treated cells (red). Quantification of average pMyosin intensity per cell (n=15 cells) and YAP nuclear to cytoplasmic ratio (n=20 cells) in shRNA treated cells. (**H**) Schematic illustrating the model of G3BP1 as the key molecular hub that forms granules at FAs enabling the function of this complex of proteins. For all graphs, significance is represented as not significant (n.s.) P > 0.05, *P ≤ 0.05, **P ≤ 0.01,***P ≤ 0.001, ****P ≤ 0.0001.

We next verify how FA-mRNAs could influence the formation of G3BP1 granules. We mixed purified G3BP1 protein with two *in vitro* transcribed FA mRNAs that mark either FA-cluster 0 or 3, and 10% of the molecular crowding agent, dextran, *in vitro*. We observed that G3BP1 formed primarily branched/elongated granules with both FA-cluster 0 and 3 mRNAs compared to when mixed with total HDF mRNA (Fig. 5D). These G3BP1 complexes were dissolved after the addition of RNase A for 5 min, confirming the key role of mRNAs in the formation of FA-mRNA G3BP1 granules *in vitro* (Fig. 5D). Overall, these data suggest that untranslated mRNAs are necessary to form G3BP1-biomolecular granules that co-localize at FAs.

To further study the contribution of G3BP1 in localizing mRNAs at FA, we used shRNAs to efficiently remove G3BP1 expression in HDFs and used smFISH to observe mRNA distribution (fig. S5D). In cells depleted for G3BP1, we found that both FA-cluster 3 and 0 markers *TRAK2* and *ZMPSTE24*, respectively, decreased their association with the respective FAs, without an overall reduction of total mRNA levels within the cells (Fig. 5E). Furthermore, HDFs depleted of G3BP1, but not G3BP2 or DDX3X, showed loss of FA properties and exhibited decreased HDF contractility associated with loss of nuclear Yap and cytosol p-myosin staining (Fig. 5F, 5G and fig. S5E). Taken together, our data suggest that G3BP1 is the key molecular component that mediate the non-coding function of mRNAs in the mechanoresponses of FAs (Fig. 5H).

Here, we showed that cytosolic mRNAs when localized to cellular organelles can form biomolecular granules with local RNA binding proteins such G3BP1, remain translationally inactive and function to organize the network composition of protein complexes such as those forming FAs. Since untranslated mRNAs do not appear to re-enter translation under conditions of FA homeostasis, depletion or activation, we speculate the FA-mRNA-G3BP1 granules have a unique structural function that is uncoupled from protein synthesis. P-bodies, so far, are the only example of cytosolic granules in normal cells that segregate translationally repressed mRNAs for decay or storage before they are reactivated for translation^48^. Interestingly, analysis of P-body proteome has shown that proteins such as G3BP1/2 are not part of these granules^49^. Therefore, we propose that our FA-mRNA-G3BP1 granules represent a new type of cytosolic mRNP where the untranslated mRNA have an unexpected non-coding role as “scaffold” molecules. Furthermore, these observations broaden the functional role of mRNA localization as a way for mRNA to exploit non-canonical function, opening the interesting possibility that this phenomenon might be conserved in other cytoplasmic organelles and regulate new cell functions.

How mRNA can function as a structural molecule is a subject for continued exploration within the field. RNAs can promote biochemical reactions directly or through helping to seed condensates by increasing the local concentration of reaction components, such as other nucleic acid molecules as well as proteins, within a particular location. For example, nuclear droplets of nascent or long-noncoding RNAs have the biophysical property to pull together even very distal regions of the genome to regulate transcriptional protein complexes^50-52^. Within the cytosol, there is evidence to suggest that localized mRNAs undergoing translation, by utilizing their 3’UTR sequence, can guide protein interactions with the nascent translated polypeptide to facilitate local signalling^8,53^. Since our FA mRNAs do not seem to be present as singletons^54^, it is possible that multiple mRNA-mRNA interactions within G3BP1 granules might help to regulate the ideal biophysical milieu for the over 200 proteins in the FA to properly organize, function, and respond to environmental cues.

Our data unexpectedly reveal that in normal somatic cells, mRNAs can mediate the interaction between FA proteins and G3BP1, a key protein required for the early condensation of stress granules. G3BP1 assemble liquid-liquid phase separation droplets by binding mRNAs not based on recognition of specific RNA sequence motifs but based on mRNA structural characteristics that include long length and the ability to form RNA-RNA interactions ^55,56^. Interestingly, we found that these RNA features are particularly enriched amongst the FA-associated mRNAs in comparison to cytosolic mRNAs, suggesting that G3BP1 may form FA mRNP granules using the same molecular principles used to form stress granules. We also found that different mRNA species localized at different FA states, which may correspond to diverse networks of FA proteins. Since RNA sequences, properties and expression levels can modulate the biophysical properties of granules, such as size, shape, surface tension, viscosity and composition^35,57^, it is possible that a more dynamic smaller FA vs a larger more stable FA could require different condensate biophysical properties to function. Since the field of biomolecular condensates is evolving rapidly, more knowledge will be required to fully understand how distinct features of mRNAs within granules can influence the structural properties of the protein complexes they are a part of.

The composition and localization of mRNAs at FAs, although never directly characterized, has been suggested since the 1990’s, where extracellular matrix promoted the formation of cytoskeletal microcompartments near newly forming FAs in fibroblasts and stained positively for ribosomal RNAs riboprobes ^58-60^. Since then, localization of β-actin mRNA and its translation at FAs was identified as an important mechanism of FA stability, cell migration and neurite outgrowth^5,61^. Our data support these initial discoveries since some FA mRNAs were found translationally active in our FA isolates, and their proteins level was accordingly regulated upon FA perturbations. However, with the exact identification and mechanistic distribution of the mRNA species at FA, our data also suggest that FA function is dependent largely on localized mRNAs with non-coding function. Inhibition of translation at FAs did not perturb FAs, although it is possible that protein lifetime might buffer the effects of blocking local translation at FA. Overall, it could be that FAs host different types of granules based on the translational status of mRNAs associated and that these complexes could share or acquire specific functions in response to different extra- or intracellular cues to modulate different cell behaviours.

Removal of RNAs or G3BP1 dissolved FA protein strata, particularly the upper-most actin-regulatory layer that is essential to transducing matrix tension into cell contractility. Cell contractility is one of the main cellular behaviours resulting from mechanotransduction of the extracellular matrix, and it is indispensable to cellular adaptation to rapid change in extracellular matrix properties. In a sense, FAs are sentinels of cellular mechanical “stress”, thus they might have developed a shared mechanism to canonical stress granules. When a cell loses the ability to sense this matrix “stress”, several pathophysiological conditions form as a result, such as wound healing defects, atherosclerosis, or cancer metastasis. The discovery that mRNAs have a non-coding function in modulating FAs open a new avenue for thinking about how aberrant matrix-cell interactions can be therapeutically corrected. The current advance in using mRNA molecules as a drug will make our data of interest for the development of future RNA therapeutics where, without affecting endogenous protein level, mRNA might regulate critical cell behaviours.

## Supplementary Materials

### Materials and Methods

#### Cell Culture and Treatments

Human umbilical vein endothelial cells (HUVECs) were purchased from Cell Applications Inc. (200-05n) and were cultured on dishes coated with 0.1% w/v gelatin (10 min at room temperature in PBS; Sigma) in M199 medium (11150-059, Thermo Fisher) supplemented with 20% FBS (F0926-500 ML, Sigma), supplemented with 10 μg/mL endothelial cell growth supplement prepared from bovine hypothalamus, 50 μg/mL heparin filtered (H3149-100KU, Sigma), and 1% Penicillin-Streptomycin (15140122, Thermo Fisher). Cells were cultured according to manufacturer’s instructions and used until passage 6. Human dermal fibroblasts (HDFs) were purchased from the American Type Culture Collection (ATCC, PCS-201–030) and cultured in fibroblast growth medium (Fibroblast Growth Kit-Low Serum; ATCC, PCS-201– 041) supplemented with 0.5 mL Phenol Red (ATCC, PCS-999-001, final concentration 33 μM) and 1% Penicillin-Streptomycin (ThermoFisher, 15140122). HDFs were cultured according to manufacturer’s recommendations and used until passage 6.

For blebbistatin treatment, cells were cultured in media supplemented blebbistatin for a final concentration of 25 uM (Sigma, B0560) for 30 minutes. The blebbistatin was diluted in DMSO, therefore control cells were treated for 30 minutes with media supplemented with an equal volume of DMSO (VWR, IC0219605525). At the end of 30 minutes, cells were fixed for analysis. For blebbistatin washout samples, at the end of the 30 minutes blebbistatin treatment, the media was removed, cells were washed, and cultured for an additional 30 minutes in media before fixation. For manganese treatment, cells were cultured in media supplemented with manganese (Sigma, M3634) for a final concentration 3 uM for 20 minutes prior to fixation. For cycloheximide (CHX) and RNase A treatments, cells were cultured in media supplemented with either CHX (Sigma, C4859-1ML) for a final concentration of 100 μg/mL or RNase A (ThermoFisher, EN0531) for a final concentration of 1 mg/ml for 10 minutes prior to either FA isolation or fixation. Extracellular matrix (ECM) coatings used were fibronectin (2 μg/cm^2^ overnight at 4 °C in PBS), laminin (1 μg/cm^2^ for 2 hours at 37°C in PBS), collagen IV (10 μg/cm^2^ overnight at 4 °C in PBS), and collagen I (10 μg/cm^2^ overnight at 4 °C in PBS).

#### Focal Adhesion Isolation

Focal adhesions were isolated as previously described ^32^. Cells were seeded on tissue culture plastic coated with bovine plasma fibronectin (10 μg/mL in PBS overnight at 4°C). After 24 hours in culture, cells were rinsed once with PBS and then treated for 3 minutes in a 2.5 mM triethanolamine (Sigma, 90279-100ML) low ionic strength buffer, pH 7.0. Immediately after the incubation, cell bodies were removed using hydrodynamic force (setting 1, Interplak, Conair) using PBS at ∼0.5 cm from and ∼90° to the surface of the dish. Dishes were immediately inspected for removal of cell bodies using phase microscopy. Isolated FAs were washed 3 times with PBS before fixation or collection from the dish using a cell scraper (LabScientific, CL-25) in the required buffer for further analysis.

#### mRNA- and Ribo-seq Library Preparation

Total RNA was extracted from four replicates of isolated FA samples (each replicate was pooled from four 100 mm dishes) or matched whole cell controls from HUVECs or HDFs using TRIzol reagent (Life Technologies) according to the manufacturer’s protocol. Prior to mRNA-seq library preparation, RNA was run on a 2100 Bioanalyzer (Agilent Technologies, Inc.) to ensure high quality RNA. Then, 500 ng of total RNA was used to prepare Lexogen QuantSeq 3′ mRNA-Seq FWD libraries for Illumina deep-sequencing according to the manufacturer’s protocols. Libraries were amplified for 13-14 PCR cycles. Ribo-Seq samples were prepared according to manufacturer’s instructions using the RiboLace Kit from Immagina BioTECHNOLOGY (RL001). For these samples, cells were treated for 5 minutes with 10 μg/mL cycloheximide (C4859-1ML, Sigma) at 37°C directly prior to focal adhesion isolation.

#### High-throughput sequencing analysis

For record keeping and configuring the bioinformatics analysis, samples annotations were stored in LabxDB ^62^. Annotations of human genes and the genome sequence of GRCh38 assembly was obtained from Ensembl 92 ^63^. The “import_ensembl” script from the FONtools ^64^ was used to (i) download FASTA and GFF3 files from Ensembl, (ii) create an index of the human genome, and (iii) generate FON1 files containing gene and transcript annotations.

#### QuantSeq RNA-seq analysis

Raw reads were first trimmed using Skewer ^65^ using the sequence AAAAAAAAAAAAAAAGATCGGAAGAGCACACGTCTGAACTCCAGTCAC with default parameters. They were then mapped to the human genome using STAR ^66^ with non-default parameters “--alignEndsType Local” and “--seedSearchStartLmaxOverLread 0.8”. FON1 files containing genes (or “metagenes”) were generated by concatenating the isoforms of each gene together using the “--method union” option of the “fon_transform” tool ^64^. Read counts per gene were computed by summing the total number of reads overlapping at least 10 nucleotides of the gene (i.e. metagene) annotation. Reads mapping to multiple loci were accounted for by dividing 1 (each read) by the number of loci to which the read was mapped to. To identify differentially expressed genes, genes with at least 1 count in both conditions in any of the replicates were first selected, and input to the “DESeq” function ^66^. The “results” function with parameters “pAdjustMethod=“fdr”, independentFiltering=FALSE” returned the differentially expressed genes.

#### Ribo-seq analysis

Raw reads were first trimmed by aligning the adaptor sequence CTGTAGGCACCATCAATAGATCGGAAGAGCACACGTCTGAACTCCAGTCAC requiring matching first five base-pairs and minimum alignment score of 60 (Matches:5, Mismatches:-4, Gap opening:-7, Gap extension:-7). Trimmed reads were then filtered of small RNAs by mapping them onto sequences extracted from RepeatMasker ^67^ (rRNA, snRNA, scRNA and tRNA types) and Ensembl ^63^ (rRNA, snoRNA, snRNA, misc_RNA, Mt_rRNA and Mt_tRNA types) using STAR ^66^ with the non-default parameters “--alignEndsType Local”, “--seedSearchStartLmaxOverLread 0.8” and “--outReadsUnmapped Fastx”. Unmapped reads (obtained from the last option above) were then mapped to human genome using STAR with the non-default options “--alignEndsType EndToEnd” and “--seedSearchStartLmaxOverLread 0.8”. The coding sequences (in genomic coordinates from FON1 file) were shifted of 12 nucleotides upstream to take into account the nucleotide separating the read start from the P-site of the translating ribosome. Read counts per mRNA were computed by summing the total number of reads overlapping at least 10 nucleotides the shifted mRNA annotation. Reads mapping to multiple loci (up to 5 loci) were accounted for by dividing 1 (each read) by the number of loci to which the read was mapped to.

#### Measurement of mRNA length, 3′UTR length and the number of AU-rich elements in 3′UTRs

mRNA length is the longest isoform of annotated protein-coding genes. 3′UTR length is the full-length 3′UTR length obtained from Refseq (hg19). For counting of AU-rich elements, we only considered the canonical sequence AUUUA. We counted the number of AU-rich elements in annotated 3′UTRs of mRNAs expressed in HDFs or HUVEC. All values are listed in Table 1.

#### Calculation of normalized ensemble diversity (NED) values

Ensemble diversity is the number of potential RNA structures that are predicted for a given RNA (Lorenz et al., 2011). As ensemble diversity increases with RNA length (Ding et al., 2005), we are using a length-normalized value (NED). RNAs with low NED values have predominantly strong local structures, whereas RNAs predicted to have high NED values often have large unstructured regions. The ensemble diversity of 3′UTR sequences was calculated using the RNAfold software (version: 2.4.14; command line: RNAfold --MEA -d2 -p -- infile=<RNA_sequences.fasta> --outfile=<RNA_sequences.RNAfold.summary>) ^68,69^ Only 3′UTRs with a length < 7500 nucleotides can be analyzed by RNAfold. To calculate NED, ensemble diversity values were divided by the length of the 3’UTR in nucleotides.

#### Probe Design

##### MERFISH probe design

The template oligo pools of Focal Adhesion mRNA MERFISH (126 transcripts) and cell cycle mRNA MERFISH (60 transcripts) were designed similarly to that previously described ^36^, for enzymatic amplification of the RNA MERFISH primary probes targeting the transcripts of focal adhesion genes and cell cycle genes, respectively. Briefly, each oligo in the template oligo pools contained six regions: 1) a 20-nt 5’ forward priming region, 2) a 20-nt readout region, 3) a 30-nt primary hybridization region, 4) a second 20-nt readout region, 5) a third 20-nt readout region, and 6) a 20-nt 3’ reverse priming region. The priming regions were selected according to a previous MERFISH study ^36^.

The 16 readout sequences were selected from a previous publication ^70^. Modified Hamming-Distance 4 (MHD4) coding scheme ^36^ was used for combinatorial barcoding. Each RNA species (gene) was represented by a unique set of 4 out of 16 readout sequences. Each oligo in the template oligo pools harbors 3 of the 4 selected readout sequences, which shuffles among the oligos targeting the same gene. Each gene is targeted by 48 oligos. Therefore, each of the 4 selected readout regions appears 36 times on the 48 oligos for the corresponding target gene.

The primary hybridization regions were chosen from the focal adhesion-enriched genes via OligoArray 2.1 ^71^ with the following criteria: 1) The melting temperature of the oligos is not lower than 66 °C; 2) the oligos do not cross-hybridize with each other at 72 °C or higher temperature; 3) the oligos cannot form stable secondary structures at 76 °C or higher temperature; 4) there is no consecutive repeats of six or more identical bases in the sequence; and 5) the oligos do not overlap with each other. The qualified oligos were then screened against human transcriptome, Ensembl 92 by BLAST+ ^72^, and oligos with significant homology with transcripts from more than one gene were excluded. 48 primary hybridization regions were selected for each target gene.

##### Single-molecule FISH probe design

The single-molecule FISH (smFISH) primary probes of the following genes were enzymatically amplified from a template oligo pool: *TLN1, CALD1, MACF1, CTNNB1, PPP1R12A, IQGAP1*. Each oligo contained four regions: 1) a 20-nt 5’ forward priming region, 2) a 30-nt readout region, 3) a 30-nt primary hybridization region, and 4) a 20-nt 3’ reverse priming region. The priming region sequences and the readout region sequence were chosen according to a previous study ^36^. The primary hybridization regions were designed as introduced above. 36 primary hybridization regions were selected for each target gene.

The smFISH primary probes of *SKP1, KPNA4, FLNB, TRAK2, KIF1C, ZMPSTE24* and *STEAP3* were individually ordered from Intergrated DNA Technology (IDT), Inc., which carried the following two regions: 1) a 30-nt primary hybridization region, and 2) a 30-nt readout region. The primary hybridization regions were designed as introduced above. 36-48 primary hybridization regions were selected for each target gene. The selected primary hybridization regions were then appended a 30-nt readout region selected from a previous study ^36^ and the whole 60-nt sequences were reverse-complemented for the order to directly yield the smFISH primary probes without the enzymatic amplification.

##### Probe synthesis

Template oligo pools were ordered from CustomArray, GenScript. MERFISH primary probes and a subset of smFISH primary probes specified in the section above were then synthesized through an enzymatic amplification procedure reported in previous studies ^36,70^. Briefly, oligo pools were amplified through limited-cycle PCR and in vitro transcription, and converted to single-stranded DNA via reverse transcription, alkaline hydrolysis and column purification. Synthesized primary probes were dried by ThermoFisher SpeedVac DNA120 Vacuum Concentrator and stored at -20 °C, and resuspended in 10 μL ultrapure water before usage. Dye-labeled readout probes, the remaining smFISH primary probes specified in the section above, and PCR primers for probe synthesis were ordered from Intergrated DNA Technology (IDT), Inc.

#### smFISH and quantification

Cells were seeded on 25 mm coverslips in 6-well dishes coated with bovine plasma fibronectin (10 μg/mL in PBS overnight at 4°C) and allowed to adhere overnight. All samples were fixed with 4% paraformaldehyde (Electron Microscopy Sciences) in PBS for 10 minutes at room temperature and permeabilized in 0.5% Triton X-100 for 10 minutes. For the IF counterstain, the incubation steps were all conducted at room temperature. Samples were first blocked in 0.3% BSA in DPBS with RNase inhibitor (1:1000, NEB M0314L) for 30 minutes, then stained with anti-PXN (1:500, Abcam, ab32084) and secondary (1:1000, ThermoFisher, A32731) antibodies in DPBS with RNase inhibitor (1:100) for 1 hour each, and finally post-fixed in 4% paraformaldehyde for 10 minutes before proceeding with the smFISH hybridization. Cells were incubated in pre-hybridization buffer (50% formamide, 2× saline-sodium citrate (SSC)) for 5 minutes at room temperature before proceeding to overnight smFISH primary hybridization. Cells were incubated at 37 °C for 16-24 h in a humid chamber in 25μL primary hybridization buffer (50% formamide, 2× SSC, 0.1% w/v yeast tRNA (Life Technologies, 15401011), 10% dextran sulfate (Sigma, D8906) and 1:100 diluted murine RNase inhibitor) containing 40nM primary probes per primary probe sequence.

After primary hybridization, cells were washed twice in 2× SSC with 0.1% v/v Tween-20 at 60 °C for 15 min each, followed by one wash in 2× SSC with 0.1% v/v Tween-20 at room temperature for 15 min. Cells were then incubated in secondary hybridization buffer (20% v/v ethylene carbonate (Sigma-Aldrich, E26258) in 2× SSC, 3nM readout probe) at room temperature for 25-30 minutes, and then washed 3 times with wash buffer (20% v/v ethylene carbonate in 2× SSC) for 5 min each. Samples mounted with Prolong Diamond Antifade with DAPI (Thermo, P36971) and imaged on a Leica SP8 or Zeiss LSM980 Airyscan 2 confocal. To quantify distance to focal adhesions, the centroid of each structure was calculated using previously published Mathematica code ^73^ and the distance between each centroid was calculated with in house Python code.

#### MERFISH imaging

MERFISH primary probe hybridization was conducted in the same way as smFISH introduced above, except that 1) cells were seeded on 1.5# coverslips of 40-mm diameter (Bioptechs, 40-1313-03193), and 2) during primary hybridization, 25uL primary hybridization buffer containing 4-8uM of primary probe sets were applied. Prior to overnight imaging, 0.1-μm yellow-green fiducial beads (Invitrogen, F8803) diluted in 2× SSC were applied to the cells, which served as markers for drift correction purpose during sequential imaging.

Imaging of the 16-bit MHD4 MERFISH scheme was conducted by 8 rounds of automated sequential hybridization, imaging and photobleaching, using a Bioptech’s FCS2 flow chamber and a home-built fluidic system ^36^. The secondary probes of MERFISH were labeled at 5’ end with either Alexa Fluor 647 or Alexa Fluor 750 dyes, and in each round of readout hybridization, two secondary probes with the two different dyes were applied simultaneously and imaged sequentially, allowing the 16-bit MHD4 codes to be read out in 8 rounds of readout hybridization.

For each round of readout hybridization, cells were first treated with hybridization buffer (20% v/v ethylene carbonate (Sigma-Aldrich, E26258) in 2× SSC containing 3nM each of the corresponding MERFISH readout probes with Alexa Fluor 647 and Alexa Fluor 750 labeling, respectively, and 1:2000 diluted murine RNase inhibitor). Cells were incubated with hybridization buffer for 20 min at room temperature, and then 2 mL of wash buffer (20% v/v ethylene carbonate in 2× SSC) was applied to remove excessive readout probes. 2mL oxygen-scavenging imaging buffer (50 mM Tris-HCl pH 8.0, 5% w/v glucose, 2 mM Trolox (Sigma-Aldrich, 238813), 0.5 mg/mL glucose oxidase (Sigma-Aldrich, G2133), 40 μg/mL catalase (Sigma-Aldrich, C30), 1:2000 diluted murine RNase inhibitor in 2× SSC) was then applied to the sample for imaging multiple fields of view. A layer of mineral oil (Sigma, 330779) was applied on top of the imaging buffer to prevent continuous oxidation.

During sequential imaging, a series of z-stack images was taken under illumination with 750-nm, 647-nm, 560-nm and 488-nm lasers sequentially. The 750-nm and 647-nm lasers were used for MERFISH readout hybridization dye excitation, the 560-nm laser for Paxillin antibody staining excitation, and the 488-nm laser for fiducial bead excitation. The step size of the z-stack images was 500 nm, with a full depth of 3.5 μm on the z dimension. The exposure time at each height was 0.4 s. After imaging was finished for all fields of view in the current readout imaging round, 2× SSC buffer was applied to the sample to allow photobleaching of MERFISH signals, which was done by exposing cells under 750-nm and 647-nm lasers for 29 s.

After 8 rounds of sequential readout hybridization and imaging, 3mL of 1:3000 diluted Hoechst (Thermo Scientific, 33342) in DPBS was applied to the sample and let stand for 2 min. Imaging buffer was then applied to the sample, and z-stack images with 405-nm laser and 488-nm laser illumination were taken as described above.

#### MERFISH imaging system

A home-built microscope was used for MERFISH experiments, which consists of a Nikon Ti2-U body, a Nikon CFI Plan Apo Lambda 60× Oil (NA1.40) objective lens, and an active auto-focusing system ^74^. A 750-nm laser (2RU-VFL-P-500-750-B1R, MPB Communications) was used to excite and image Alexa Fluor 750 on readout probes. A 647-nm laser (2RU-VFL-P-1000-647-B1R, MPB Communications) was used to excite and image Alexa Fluor 647 on readout probes. A 560-nm laser (2RU-VFL-P-1000-560-B1R, MPB Communications) was used to excite and image the secondary antibody for Paxillin staining. A 488-nm laser (2RU-VFL-P-500-488-B1R, MPB Communications) was used to excite and image the yellow-green fiducial beads for drift correction. A 405-nm laser (OBIS 405 nm LX 50 mW, Coherent) was used to excite and image the Hoechst stain. A multi-band dichroic mirror (ZT405/488/561/647/752rpc-UF2, Chroma) was used on the excitation path to direct the five laser lines to the sample. A multi-band emission filter (ZET405/488/561/647-656/752 m, Chroma) and a Hamamatsu Orca Flash 4.0 V3 camera were installed on the emission path. The pixel size of the imaging system was 108 nm. An automated motorized x-y sample stage (SCAN IM 112×74, Marzhauser) was used to automatically image multiple fields of view.

#### MERFISH analysis

##### MERFISH decoding

MERFISH images were processed using MATLAB version R2020a using a previously reported pipeline ^75^. Briefly, sample drift between sequential imaging rounds was corrected by fitting the fiducial beads markers with 2D Gaussian functions, identifying the movement of their center positions in x-y plane between the current imaging round and the first imaging round, and subtracting the movement by image translation. After drift correction, the background of each image was derived by image opening using a disk-shaped structural element with an empirically determined radius of 5 pixels, and the background were subtracted from the raw RNA MERFISH image. The potential RNA signals were then identified by finding the regional maxima with pixels above a threshold, binarization, and dilation using a square structural element with a width of 3 pixels. To determine the optimal threshold for identifying RNA signals, an adaptive thresholding procedure was applied to each round of imaging, so that the relative abundance of identified RNA molecules in the imaging rounds fit the expected relative abundance derived from bulk RNA-seq data. This process was applied to all MERFISH readout imaging rounds at all z-stacks for all fields of views. Afterwards, at each z height, all 16-bits of binarized MERFISH readout images were assembled, and a 16-bit binary code was determined for each pixel. Pixels with a binary code that existed in the MERFISH codebook were recognized as RNA molecules. Adjacent pixels sharing the same code were processed as the same RNA molecule. For downstream analysis, the identified RNA molecules at each height were projected onto a single x-y plane along the z dimension.

##### Cell segmentation

Cells were segmented based on dilated MERFISH signals and Hoechst nuclear staining using a watershed-based algorithm. First, the movement between the fiducial beads in the Hoechst imaging round and the fiducial beads in the first MERFISH imaging round was subtracted as mentioned above. In some datasets, uneven illumination was observed in the 405-nm channel, and in such cases the illumination was flattened by adaptive thresholding with an empirically determined neighborhood size of 401 pixels. Hoechst nuclear staining was max-projected along the z dimension, re-scaled to a 0-1 grayscale image, and the image intensity was adjusted to so that the top 1% and the bottom 1% pixels were saturated. The Hoechst images then underwent image opening and closing by reconstruction with a disk-shaped structural element with a radius of 20 pixels to serve as foreground markers during watershed segmentation. The mean-projection of raw RNA FISH signal from both 647-nm channel and 750-nm channel in the first hybridization round was merged and opened-closed by reconstruction using a disk-shaped structural element with a radius of 20 pixels, which served as the background marker.

##### Focal adhesion pattern extraction

The Paxillin antibody staining in the 560-nm channel of the first imaging round was used to extract focal adhesion patterns. The z-stack of Paxillin antibody staining underwent max projection along the z dimension. Top-hat filtering was then performed on the max-z projected images with a disk-shaped structural element with a radius of 4 or 6 pixels. The intensity of the filtered images was then re-scaled and adjusted as mentioned above, after which the images were binarized. Focal adhesion patterns were extracted from the binarized images by removing small objects from the binarized image that were smaller than 90 or 100 pixels. The mentioned parameters that had variable values were manually optimized for each individual dataset to account for minor batch differences in labeling intensity so that the binary focal adhesion patterns best resembled those in the raw images. We noticed that high antibody staining background tended to appear in peri-nuclear region occasionally, which could be mis-identified as real focal adhesions, and therefore focal adhesion patterns within 15 pixels to the nucleus edge were removed.

##### Distance measurement among focal adhesion, RNA and nucleus

The RNA molecules were z-projected onto the x-y plane to calculate the 2D distance between RNAs and other cellular structures. The edges of binarized focal adhesions and nuclei were extracted by Sobel methods, and the positions of the points on the edges were stored and further used in measuring distance between RNA and focal adhesion edge. The positions of focal adhesion centroids and points on the nuclear edges were used to measure distance between focal adhesion and nuclear edge.

##### Focal adhesion clustering and grouped focal adhesion property analysis (PHATE)

To generate RNA profile for each focal adhesion from MERFISH images, focal adhesion patterns were extracted as mentioned above, and a focal adhesion-adjacent region was defined by dilating the focal adhesion patterns with a disk-shaped structural element with a radius of 20 pixels. The RNAs that resided in the focal adhesion or the focal adhesion-adjacent region in a 2D projection were considered to “belong to” the current focal adhesion. Each focal adhesion was also assigned a unique index. A matrix was generated for each MERFISH dataset, with each row representing a focal adhesion, and each column representing the count of RNA molecules that were considered to belong to the current focal adhesion for each RNA species. Five such matrices generated from five MERFISH replicates were then imported into the Seurat package ^76^ in R and merged.

The focal adhesions were first filtered according to the number RNA species they exhibited, and only those with more than 3 non-translated mRNA species were retained. The excluded focal adhesions were assigned as “clusterNA”. Then the RNA profiles of each single focal adhesion were normalized and scaled using standard parameters, and principle component analysis (PCA) was performed. Louvain clustering was subsequently applied on the first six principle components with a resolution of 0.3, and the markers for each cluster was determined with the following criteria: 1) only genes that were detected in a minimal percentage of 25% focal adhesions of one cluster were considered (min.pct = 0.25); and 2) only genes that showed on average at least 0.25-log-fold difference between two clusters were considered (logfc.threshold = 0.25). To visualize the grouping of identified clusters, PHATE package ^40^ was applied with a number of 10 principle components to find neighbors. Feature plots of selected RNA species were plotted using a minimal cutoff of 9^th^ quantile (min.cutoff = “q9”).

To compare the size, circularity and the distance to nuclear edge, focal adhesions of different clusters were then recovered in the images based on the unique focal adhesion indices, and the measurements were taken. Specifically, the size and circularity of focal adhesions were calculated using the regionprops function in MATLAB, and the size in pixel was subsequently converted to that in μm2. The statistical analysis was performed in GraphPad Prism using Kruskal-Wallis test, and pairwise comparisons were conducted using Dunn’s statistical hypothesis testing with alpha = 0.05.

#### Mass spectrometry sample preparation and analysis

For proteomics analysis, total protein was extracted from isolated FA samples or matched whole cell controls from HUVECs or HDFs using TRIzol reagent (Life Technologies) according to the manufacturer’s protocol. Similarly, TRIzol was used on isolated FAs from control cells, cells treated with cycloheximide, or cells treated with RNase A (four replicates for each group). For all samples, the dried protein was sent to the Keck Mass Spectrometry & Proteomics Resource of the W.M. Keck Foundation Biotechnology Resource Laboratory at Yale University. Protein abundances were normalized to total abundances of detected spectral peaks for each sample.

#### Puro-PLA and quantification

Puro-PLA protocols were conducted as described previously ^77^. Here, cells were seeded on 25 mm coverslips in 6-well dishes coated with bovine plasma fibronectin (10 μg/mL in PBS overnight at 4°C) and allowed to adhere overnight. Samples were treated with 1 uM Puromycin (Thermo, A1113803) for 5 minutes then fixed in 4% paraformaldehyde (Electron Microscopy Sciences) in PBS and permeabilized in 0.5% Triton X-100 for 5 minutes. Proximity ligation was performed using the Duolink kit from Millipore Sigma (DUO92101) according to manufactures instructions. Primary antibodies used for Puro-PLA were specific to puromycin (1:5000, Millipore Sigma, MABE343), KIF1C (1:50, Abcam, ab125903), TRAK2 (1:100, Sigma, HPA062163), CTNNB1 (1:1000, Sigma, C2206), NET1 (1:500, Abcam, ab113202), TLN1 (1:500, Abcam, ab71333), FLNA (1:500) ^78-81^. All samples were counterstained with anti-PXN primary (1:40, R&D Systems, AF4259) and secondary (1:1000, ThermoFisher, A-11015) antibodies to identify focal adhesions. Samples were mounted with Duolink Mounting Media with DAPI (Millipore Sigma, DUO8204) and imaged on a Leica SP8 confocal. To quantify distance to focal adhesions, the centroid of each structure was calculated using previously published Mathematica code^73^ and the distance between each centroid was calculated with in house Python code.

#### Cell immunostaining and quantification

For immunofluorescence analysis, FAs were fixed with 4% paraformaldehyde (Electron Microscopy Sciences) in PBS for 10 minutes. Cells were then washed and permeabilized in 0.5% Triton X-100 in DPBS with calcium and magnesium (Thermo, 21300025) for 5 minutes. Cells were blocked with 10% serum for 1 hour, incubated for either 1 hour at room temperature or overnight at 4°C with primary antibodies, and incubated for 1 hour with secondary antibodies. Primary antibodies used were specific to VCL (1:500, Sigma, V9131), PXN (1:500, Abcam, ab32084), ITGB1 (1:100, Abcam, ab30394), TLN1 (1:500, Abcam, ab71333), ACTN1 (1:100, Santa Cruz, sc-17829), YAP (1:200, Santa Cruz, sc-101199), p-MCL2 (pmyosin, 1:400, Cell Signaling, 3671), G3BP1 (1:400, Santa Cruz, sc-365338), G3BP2 (1:400, Sigma, HPA018425), and DDX3X (1:200, Bethyl Labs, A300-474A). Secondary antibodies for either rabbit (1:1000, ThermoFisher, A-32731, A-21244) or mouse (1:1000, ThermoFisher, A-11001 or A-21235). Samples were mounted with DAPI in Fluoromount-G (SouthernBiotech) and imaged on a Leica SP8 confocal or Zeiss LSM980 Airyscan 2 confocal. To quantify properties of focal adhesions (size, circularity, number, distance to nuclei) previously published Mathematica code was used^73^. To quantify the intensity of FA proteins, the plot profile function in Image J was used. A region of interest was drawn across the length of the FA and the intensity of each protein was quantified and normalized to the max value. To align the FA proteins, the length of each FA was normalized to 1 and PXN or VCL were used as reference structures. YAP staining was quantified by taking the average nuclear YAP signal (overlapping with the DAPI stain) and dividing by the average cytoplasmic YAP signal (area in the non-nuclear cell mask). pMyosin signal was quantified by taking the average signal within the cell. For G3BP1 granules, images were thresholded in Image J and the analyze particles function was used to quantify their size.

#### Western Blots

For western blot analysis, cells or FA isolates were collected in RIPA buffer (Thermo 89900) with protease inhibitors (1:200, Millipore 539134) and protein concentration was determined by BCA assay (Thermo, 23225). Samples were loaded onto 4–12% SDS–PAGE gels, transferred on a Immun-Blot PVDF membrane (Bio-Rad), and blocked with 5% BSA or 5% milk for 1 hour. Blots were then incubated with primary antibodies overnight at 4 °C. Primary antibodies used were specific to VCL (1:5000, Sigma, V9131), PXN (1:8000, Abcam, ab32084), TLN1 (1:2500, Abcam, ab104913), GAPDH (1:8000, Cell Signaling, 2118), AKT1/2/3 (1:200, Santa Cruz, sc-81434), α-Tubulin (1:1000, Millipore, 05-829), β-actin (1:500, Santa Cruz, sc-130656), PDI (1:400, Santa Cruz, sc-474551), TOM20 (1:100, Santa Cruz, sc-17764), G3BP1 (1:500, Santa Cruz, sc-365338), G3BP2 (1:1000, Sigma, HPA018425), and DDX3X (1:2500, Bethyl Labs, A300-474A). Then, membranes were washed and incubated with secondary antibody anti-rabbit-HRP or anti-mouse-HRP (1:8000, Cell Signaling, 7076 and 7074) for 1 h at room temperature. After washing, blots were developed with super signal west pico chemiluminescent substrate (Thermo) using a SYNGENE G-Box imager. Quantification was performed using ImageJ.

#### shRNA knockdown of G3BP1, G3BP2, or DDX3X

Knockdown of RNA binding proteins was performed using MISSION shRNA Plasmid DNA Vectors (Sigma-Aldrich). Lentiviruses were prepared in Lenti-X 293T cells using shRNAs targeted to G3BP1 (TRCN0000008723), G3BP2 (TRCN0000047550), DDX3X (TRCN0000000002), or a non-specific control (pLKO.1-puro non-Mammalian shRNA Control Plasmid DNA, SHC002). Briefly, shRNA plasmids (3 μg) were mixed with packaging (psPAX2, 2.25 μg) and envelope (pMD2.G, 750 ng) plasmids, then mixed with Lipofectamine 2000 (Thermo Fisher) for 20 minutes before being added to cells. Virus containing supernatant was collected at 36 and 60 h post-transfection, filtered, and stored at -80°C. Cells were then infected with filtered virus in the presence of polybrene (8 μg/ml) and used for experiments 3-5 days post infection.

#### G3BP1 *in vitro* phase separation assay

Phase separation assay experiments were all conducted at room temperature using human G3BP1 protein (NBP1-50925, Novus Biologicals). Whole cell RNA was extracted from HUVECs using TRIzol reagent (Life Technologies). Cluster 0 and cluster 3 RNA was *in vitro* transcribed using the T7 MEGAscript kit (AM1334, Thermo Fisher) according to the manufacturer’s protocol. Template sequences were designed and purchased using GeneArt Gene synthesis from Invitrogen and a T7 promoter was added to the 5’ end of full-length mRNA sequences for IVT. The transcription reaction was incubated overnight at 37°C and RNA was purified using phenol chloroform extraction and isopropanol precipitation. RNAs were dissolved in Nuclease-free water and stored at -20°C. Prior to LLPS, a nanodrop was used to determine the concentration of RNAs and they were diluted accordingly. All experiments were conducted with 100 ng/μl RNA, 5 μM G3BP1, and 10% dextran as a crowding agent. RNA and protein were mixed in PCR tubes before being transferred to a chamber created by a coverslip, a glass slide, and a double-sided spacer (Sigma GBL654002). Samples were imaged using a DIC microscope using a Nikon 80i microscope with a 20x DIC objective. All imaged were captured within 20 min after LLPS induction.

#### Statistics

All the statistical analysis was performed using Prism version 9.2.0. Unless otherwise noticed, values were analyzed with either *t*-tests or one-way analysis of variance (ANOVA), with pair-wise comparisons using a Tukey test as appropriate. Figure legends indicate the exact number of measurements and samples for each test.

**Fig S1.**
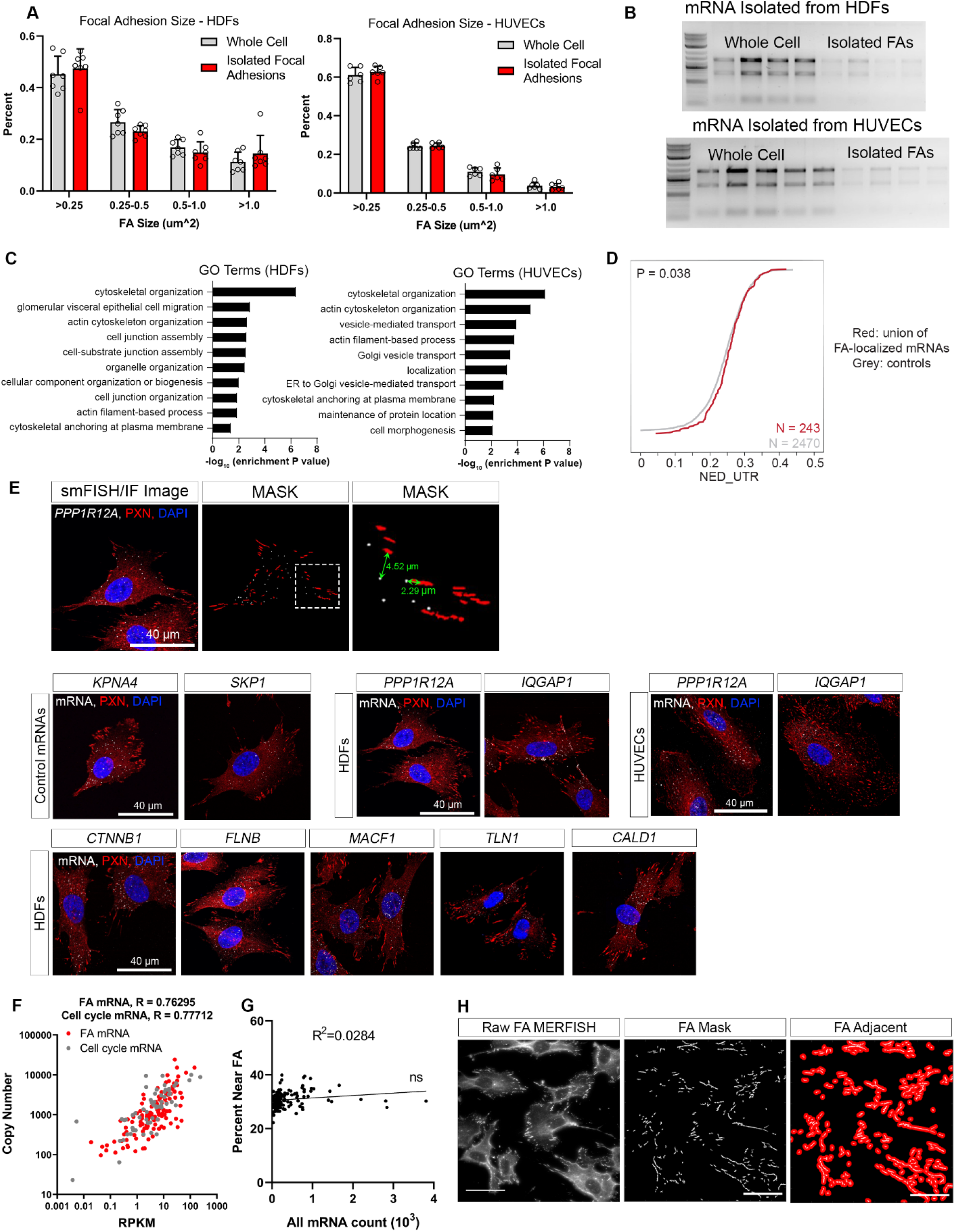
Validation of FA isolation and FISH quantification. (**A**) Histogram of the size distribution of vinculin (VCL)-containing FAs in images of whole cell or isolated FAs (Mean ± STDEV for each bar, n=7 cells). No significant difference was detected in the percentage of FAs of each size range between the two groups in either HDFs or HUVECs. (**B**) Purified RNA from whole cell and isolated FA samples in both HDFs and HUVECs. RNA was further prepared for mRNA sequencing. (**C**) Top 10 GO terms for localized mRNAs in HDFs and HUVECs sorted by their p-value. Cytoskeletal organization was the top GO term in both cell type. (**D**) NED score, a representation of structure and intermolecular interactions, for the 3’UTR of mRNAs. Cumulative histogram plot of NED values in the 3’UTR for control mRNAs (grey) and FA localized mRNA (red). (**E**) (Top) Method for quantifying distance from mRNA to nearest FA. A threshold mask was applied to both the smFISH and Paxillin (FA, PXN) images to identify mRNA (white) and PXN (red). The centroid of each structure was calculated using previously published Mathematica code^73^ and the distance between each centroid was calculated with in house Python code. Data is represented as the distance from one mRNA molecule to its nearest FA. (Bottom) representative images of each mRNA species (white) and FA (PXN, red). Control mRNAs (*KPNA4, SKP1*) and localized mRNAs (*PPP1R12A, IQGAP1, CTNNB1, FLNB, MACF1, TLN1, CALD1*) were tested. (**F**) Scatter plot of the average copy number of each RNA species from MERFISH versus the abundance determined by bulk sequencing in RPKM. Values are shown for both FA mRNA MERFISH library (R=0.74336) and the Cell Cycle MERFISH library (R=0.77712). (**G**) Abundance of mRNA vs the percent mRNA near FA (defined at >2.2 μm from FA center) from MERFISH data. Linear regression analysis found the slop was not significantly non-zero with R^2^=0.0284. (**H**) Raw FA images from MERFISH, the FA mask applied after MERFISH imaging, and the FA adjacent pixels show in red. For all graphs, significance is represented as **P ≤ 0.01,***P ≤ 0.001.

**Fig S2.**
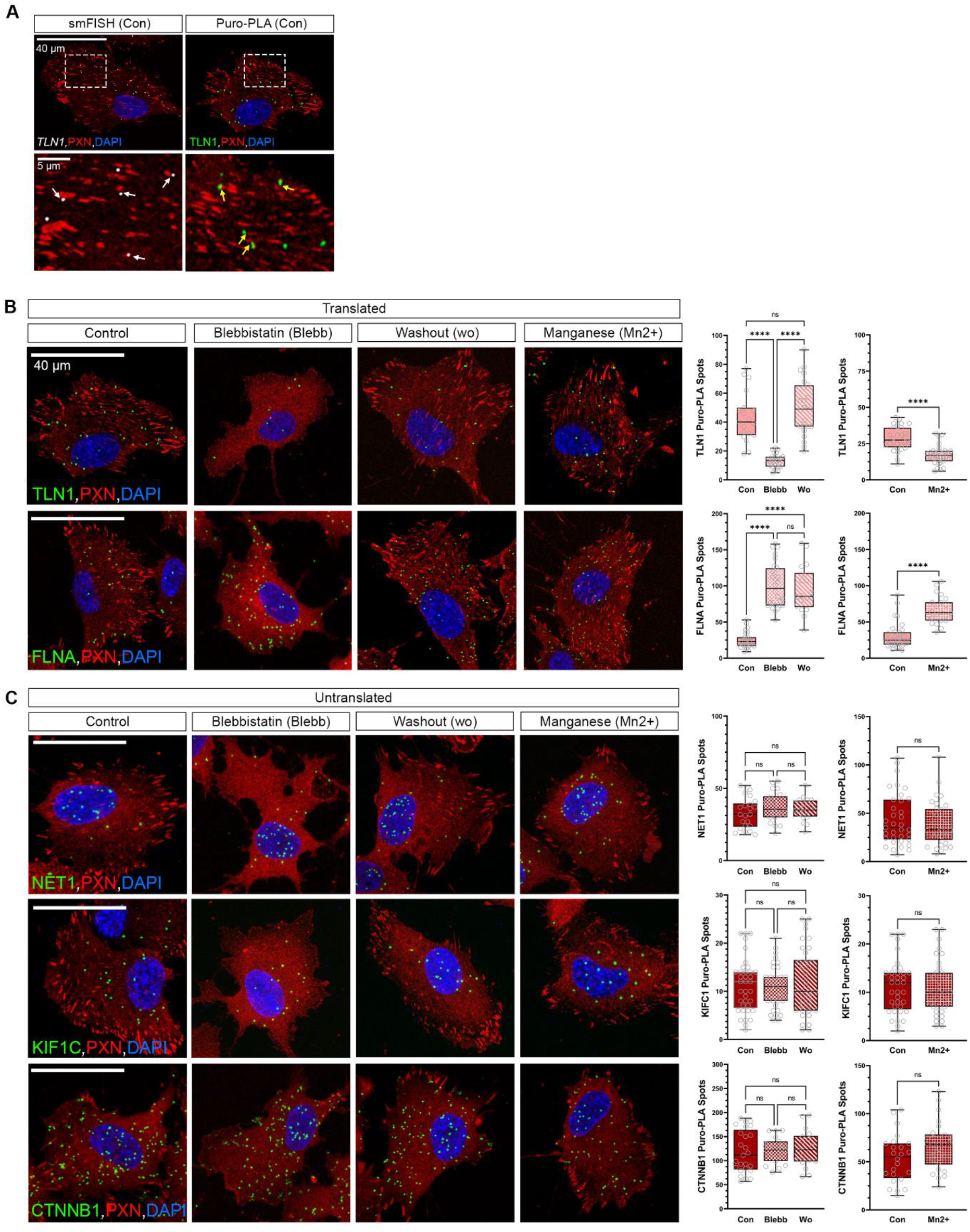
Translation in cells with FA perturbation. (**A**) *TRAK2* mRNA (white) and spots of TRAK2 translation (green) in HDFs counterstained for FA protein (PXN, red) and DAPI (blue). Both the mRNA and the translational spots are found near FAs with areas of localization indicated by arrows (white for mRNA, yellow for Puro-PLA). Representative Puro-PLA images for translated (**B**) (TLN1, FLNA) and untranslated (**C**) (NET1, KIF1C, CTNNB1) mRNA in control, blebbistatin treated (25 uM for 30 minutes), washout (30 minutes after removal of blebbistatin), or manganese treated cells (3 uM for 20 minutes). All spots of translation are show in green and cells are counterstained with FA (VCL, red) and DAPI (blue). Quantification shown as box plots for the overall number of translational events per cell (n=20-30 cells). For all graphs, significance is represented as not significant (n.s.) P > 0.05, ****P ≤ 0.0001.

**Fig S3.**
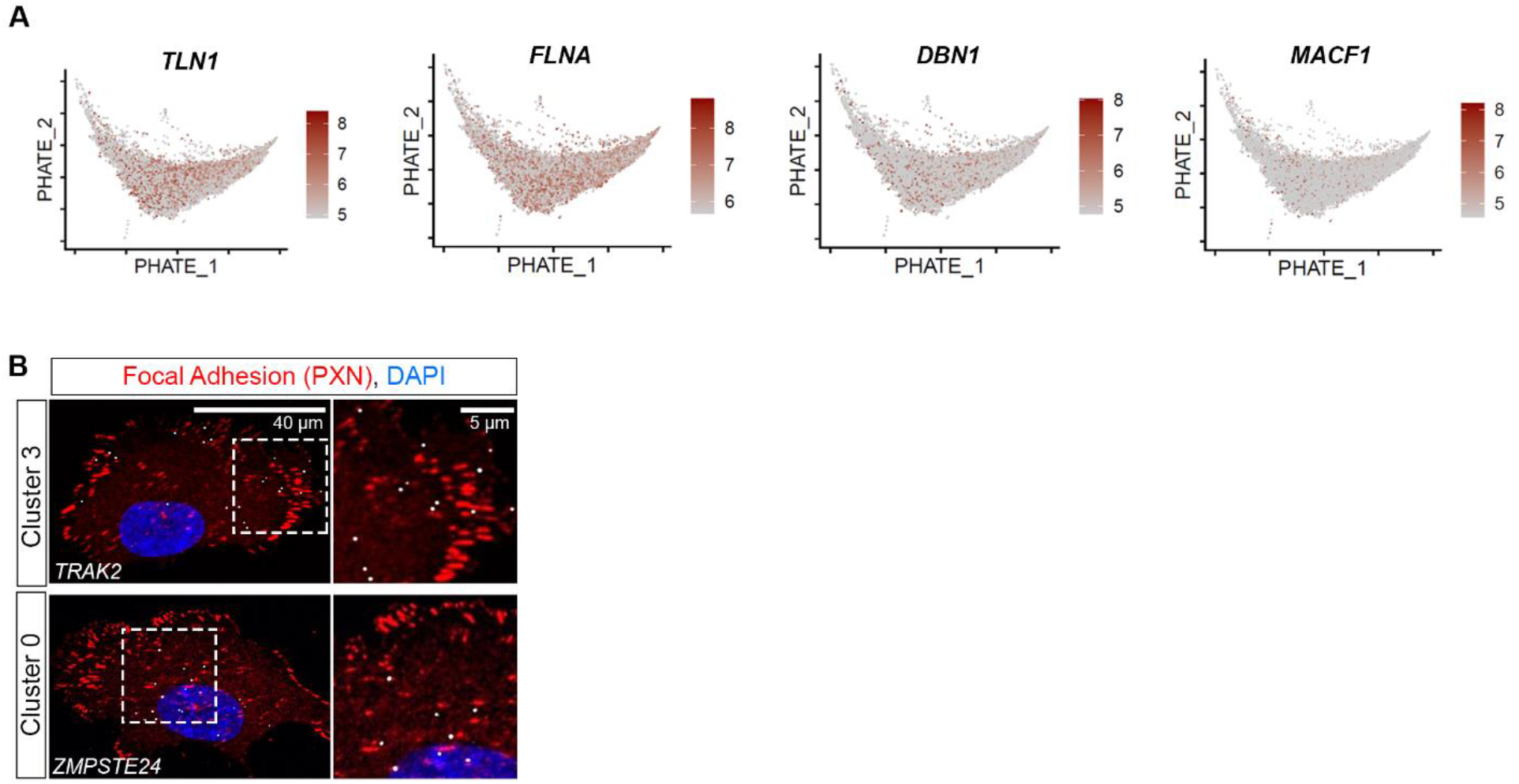
Specific mRNA species mark FA clusters. (**A**) Feature plots of three translated mRNA species (*TLN1, FLNA, DBN1, MACF1*). Expression is distributed throughout the four FA clusters. (**B**) Representative images of cluster 3 (*TRAK2*) or cluster 0 (*ZMPSTE24*) mRNA (white), FA (PXN, red), and DAPI. Doted boxes depicted on left panels represent the region highly magnified in the right panels.

**Fig S4.**
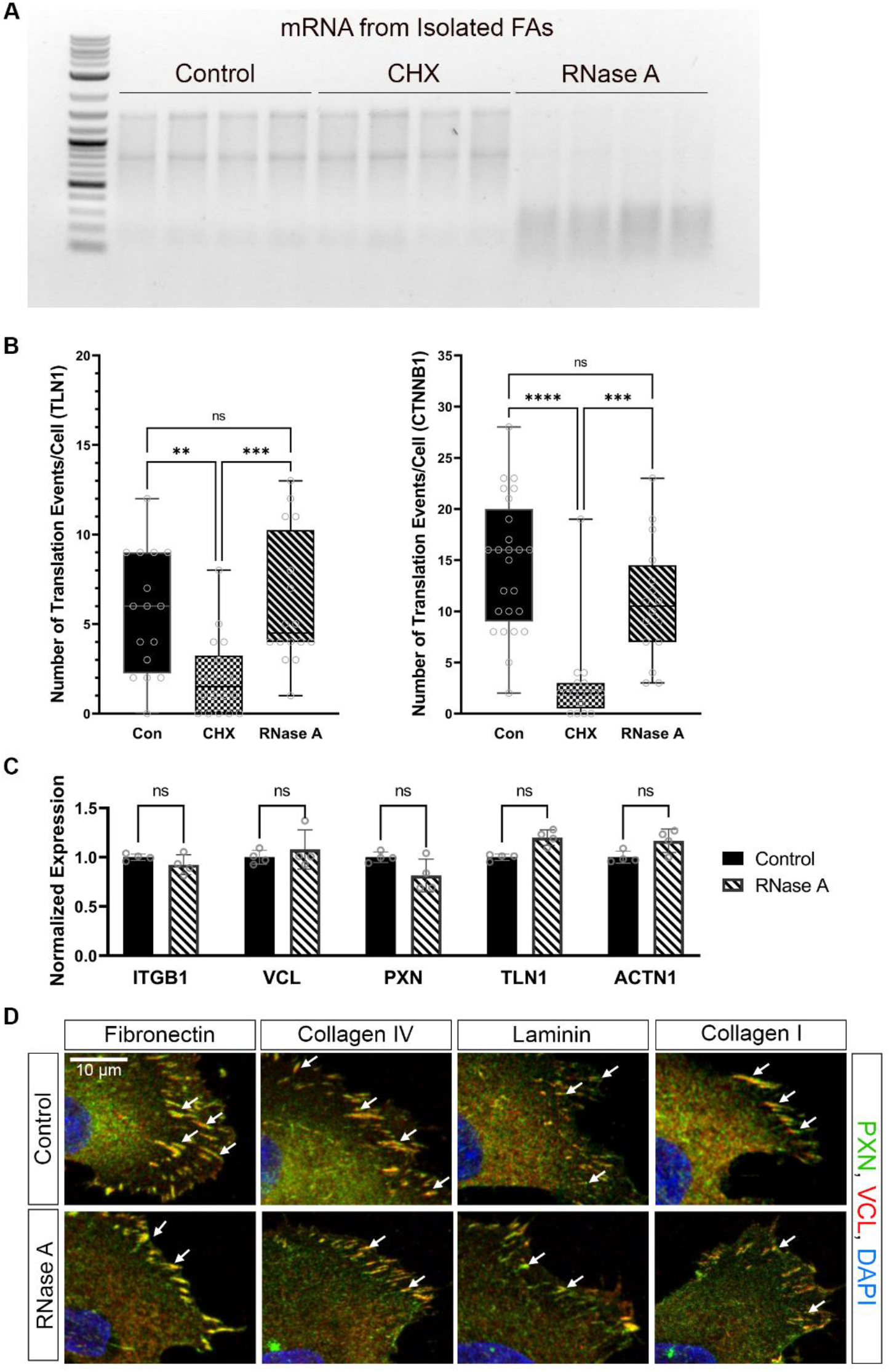
Analysis of RNase A treatment. (**A**) RNA purified using standard protocols using Trizol from isolated FAs from control, cyclohexamide (CHX, 100 μg/mL for 10 minutes), or RNase A treatment (1 mg/ml for 10 minutes). (**B**) Box plots represent the number of translational events detected per cell using Puro-PLA for TLN1 and CTNNB1 in Con, CHX, and RNase A treated cells (n=25 cells). CHX treatment decreased the amount of newly synthesized peptide while RNase A treatment saw no change compared to control cells. (**C**) Mean ± STDEV expression of core FA proteins (ITGB1, VCL, PXN, TLN1, ACTN1) from liquid chromatography–mass spectrometry analysis of isolated FAs from control and RNase A treated cells (normalized to controls). (**D**) Representative images of either control or RNase A (1 mg/ml for 10 minutes) treated cells cultured on fibronectin (FN), collagen IV (Col IV), laminin (LM), or collagen I (col I). FAs (white arrows) marked by paxillin (PXN, green) and Vinculin (VCL, red) are show with DAPI (blue) counterstain. For all graphs, significance is represented as not significant (n.s.) P > 0.05, **P ≤ 0.01,***P ≤ 0.001, ****P ≤ 0.0001.

**Fig S5.**
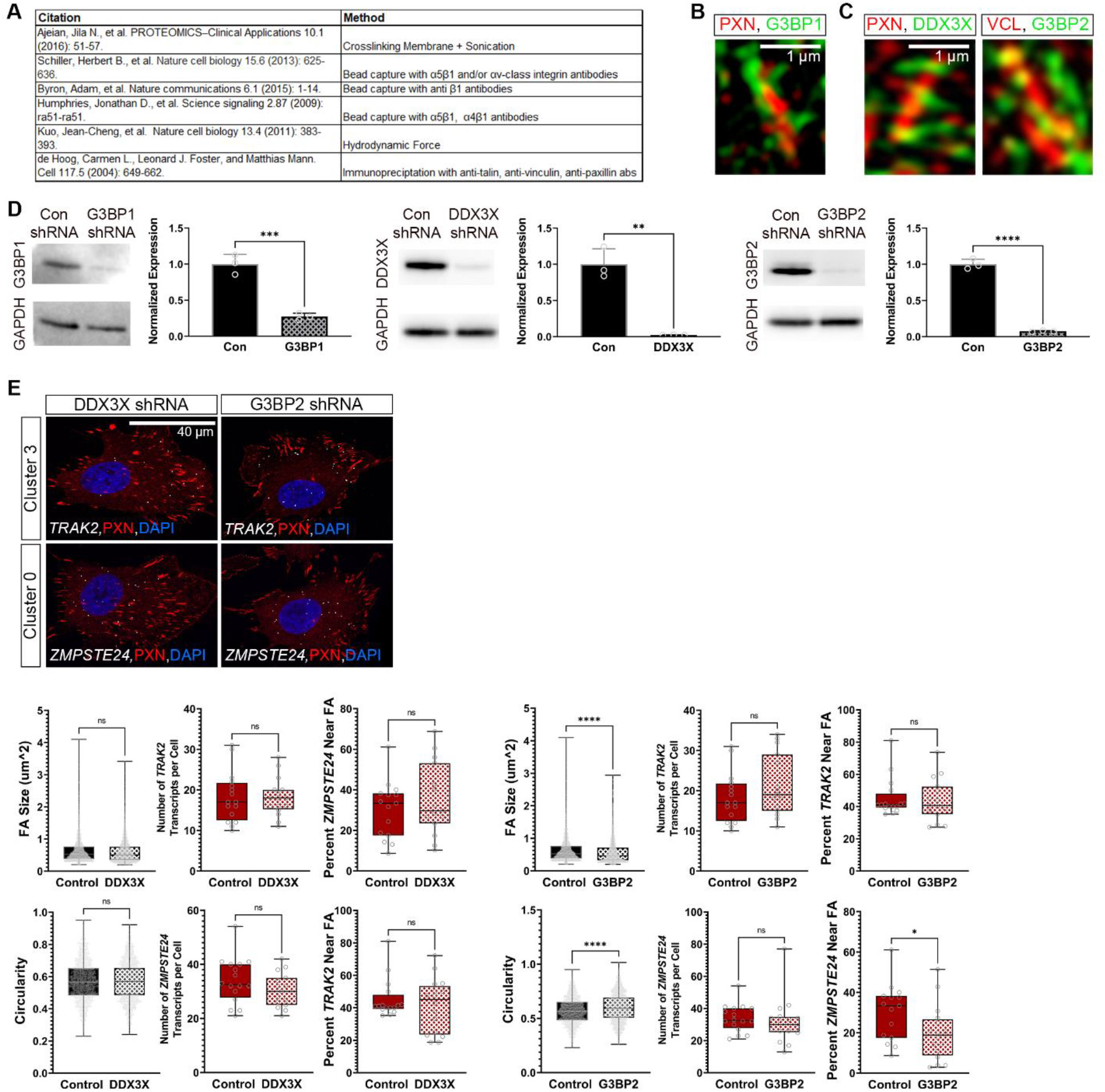
Effect of mRNPs on FA morphology. (**A**) Papers and their corresponding methods that identified G3BP1 in FAs. (**B**) Representative image of FA (PXN, red), G3BP1 (green) and DAPI for FA in cluster 0. (**C**) Representative image for RBPs (DDX3X or G3BP2, green) and FA (PXN, red) taken with high resolution imaging. (**D**) Western blot images and quantification (Mean ± STDEV) of G3BP1, DDX3X, and G3BP2 after knockdown with shRNA delivered via lentivirus (4 days post infection). Quantification is normalized to GAPDH (n=3 independent knockdowns). Control (con) samples were treated with a non-specific shRNA. (**E**) Representative images of mRNAs (cluster 3, *TRAK2*; cluster 0, *ZMPSTE24*) and FAs (PXN, red) in cells treated with shRNAs against DDX3X or G3BP2. Box plots of FA size and FA circularity, as well as the number of mRNA transcripts per cell or the percent mRNA near FA (defined at >2.2 μm from FA center) for each cluster marker. For all graphs, significance is represented as not significant (n.s.) P > 0.05, *P ≤ 0.05, **P ≤ 0.01,***P ≤ 0.001, ****P ≤ 0.0001.

**Table S1. Table of mRNAs and their properties enriched in focal adhesion samples.** The most abundant transcript and its corresponding Gene ID for mRNAs identified in focal adhesion (FA) samples from human dermal fibroblasts (HDFs), human umbilical vein endothelial cells (HUVECs), or in both cell types. The properties, including NED, overall length, and 3’UTR length, of transcripts and control mRNAs, not statistically enriched in FA samples.

**Table S2. MERFISH and smFISH probe sequences, primers, and codebook.** The codebook for FA RNA and cell cycle MERFISH used to decode the images. Primers to generate MERFISH library and individual smFISH probes from their given probe sequences. Secondary sequences and their conjugated fluorophores for FISH imaging. Some of the smFISH probes were purchased directly from IDT and the secondary probe sequence: 5’ dye label - GGGCGTATAACCGTCGCCACGCGGACGCAA-3’ with the 5’ Alexa Fluor^®^ 647 (NHS Ester) conjugation.

**Table S3. Translating mRNA species localized to focal adhesions.** The most abundant transcript and its corresponding Gene ID for mRNAs identified in focal adhesion (FA) samples that were also identified in the Ribo-seq and proteomics analysis. mRNAs are separated by cell type, HDF and HUVEC, or in both.

**Table S4. Proteomics analysis from RNase A and CHX treatment.** Normalized counts from mass spectrometry analysis of FA isolates from control, CHX, or RNase A treated cells. For all samples N=4 independent replicates. Values within the table are described in detail within the first worksheet tab.

## References

1 Holt, C. E. & Bullock, S. L. Subcellular mRNA localization in animal cells and why it matters. Science 326, 1212–1216, doi:10.1126/science.1176488 (2009).

2 Wang, E. T. et al. Dysregulation of mRNA Localization and Translation in Genetic Disease. J Neurosci 36, 11418–11426, doi:10.1523/JNEUROSCI.2352-16.2016 (2016).

3 Fernandopulle, M. S., Lippincott-Schwartz, J. & Ward, M. E. RNA transport and local translation in neurodevelopmental and neurodegenerative disease. Nat Neurosci 24, 622–632, doi:10.1038/s41593-020-00785-2 (2021).

4 Rongo, C., Gavis, E. R. & Lehmann, R. Localization of oskar RNA regulates oskar translation and requires Oskar protein. Development 121, 2737–2746 (1995).

5 Katz, Z. B. et al. beta-Actin mRNA compartmentalization enhances focal adhesion stability and directs cell migration. Genes Dev 26, 1885–1890, doi:10.1101/gad.190413.112 (2012).

6 Banani, S. F., Lee, H. O., Hyman, A. A. & Rosen, M. K. Biomolecular condensates: organizers of cellular biochemistry. Nat Rev Mol Cell Biol 18, 285–298, doi:10.1038/nrm.2017.7 (2017).

7 Benhalevy, D., Anastasakis, D. G. & Hafner, M. Proximity-CLIP provides a snapshot of protein-occupied RNA elements in subcellular compartments. Nat Methods 15, 1074–1082, doi:10.1038/s41592-018-0220-y (2018).

8 Lee, S. H. & Mayr, C. Gain of Additional BIRC3 Protein Functions through 3’-UTR-Mediated Protein Complex Formation. Mol Cell 74, 701–712 e709, doi:10.1016/j.molcel.2019.03.006 (2019).

9 Jung, H., Gkogkas, C. G., Sonenberg, N. & Holt, C. E. Remote control of gene function by local translation. Cell 157, 26–40, doi:10.1016/j.cell.2014.03.005 (2014).

10 Wang, Y. et al. Genome-wide screening of NEAT1 regulators reveals cross-regulation between paraspeckles and mitochondria. Nat Cell Biol 20, 1145–1158, doi:10.1038/s41556-018-0204-2 (2018).

11 Protter, D. S. W. & Parker, R. Principles and Properties of Stress Granules. Trends Cell Biol 26, 668–679, doi:10.1016/j.tcb.2016.05.004 (2016).

12 Decker, C. J. & Parker, R. P-bodies and stress granules: possible roles in the control of translation and mRNA degradation. Cold Spring Harb Perspect Biol 4, a012286, doi:10.1101/cshperspect.a012286 (2012).

13 Humphrey, J. D., Dufresne, E. R. & Schwartz, M. A. Mechanotransduction and extracellular matrix homeostasis. Nat Rev Mol Cell Biol 15, 802–812, doi:10.1038/nrm3896 (2014).

14 Mazzag, B. & Barakat, A. I. The effect of noisy flow on endothelial cell mechanotransduction: a computational study. Ann Biomed Eng 39, 911–921, doi:10.1007/s10439-010-0181-5 (2011).

15 Wynn, T. A. & Ramalingam, T. R. Mechanisms of fibrosis: therapeutic translation for fibrotic disease. Nat Med 18, 1028–1040, doi:10.1038/nm.2807 (2012).

16 Bangasser, B. L. et al. Shifting the optimal stiffness for cell migration. Nat Commun 8, 15313, doi:10.1038/ncomms15313 (2017).

17 Marquis, M. E. et al. Bone cells-biomaterials interactions. Front Biosci (Landmark Ed) 14, 1023–1067, doi:10.2741/3293 (2009).

18 Hu, Y. L. et al. FAK and paxillin dynamics at focal adhesions in the protrusions of migrating cells. Sci Rep 4, 6024, doi:10.1038/srep06024 (2014).

19 Schwartz, M. A. Integrins and extracellular matrix in mechanotransduction. Cold Spring Harb Perspect Biol 2, a005066, doi:10.1101/cshperspect.a005066 (2010).

20 Parsons, J. T., Horwitz, A. R. & Schwartz, M. A. Cell adhesion: integrating cytoskeletal dynamics and cellular tension. Nat Rev Mol Cell Biol 11, 633–643, doi:10.1038/nrm2957 (2010).

21 Ajeian, J. N. et al. Proteomic analysis of integrin-associated complexes from mesenchymal stem cells. Proteomics Clin Appl 10, 51–57, doi:10.1002/prca.201500033 (2016).

22 Schiller, H. B. et al. beta1-and alphav-class integrins cooperate to regulate myosin II during rigidity sensing of fibronectin-based microenvironments. Nat Cell Biol 15, 625–636, doi:10.1038/ncb2747 (2013).

23 Byron, A. et al. A proteomic approach reveals integrin activation state-dependent control of microtubule cortical targeting. Nat Commun 6, 6135, doi:10.1038/ncomms7135 (2015).

24 Humphries, J. D. et al. Proteomic analysis of integrin-associated complexes identifies RCC2 as a dual regulator of Rac1 and Arf6. Sci Signal 2, ra51, doi:10.1126/scisignal.2000396 (2009).

25 Kuo, J. C., Han, X., Hsiao, C. T., Yates, J. R., 3rd & Waterman, C. M. Analysis of the myosin-II-responsive focal adhesion proteome reveals a role for beta-Pix in negative regulation of focal adhesion maturation. Nat Cell Biol 13, 383–393, doi:10.1038/ncb2216 (2011).

26 de Hoog, C. L., Foster, L. J. & Mann, M. RNA and RNA binding proteins participate in early stages of cell spreading through spreading initiation centers. Cell 117, 649–662 (2004).

27 Zappulo, A. et al. RNA localization is a key determinant of neurite-enriched proteome. Nat Commun 8, 583, doi:10.1038/s41467-017-00690-6 (2017).

28 Moissoglu, K., Yasuda, K., Wang, T., Chrisafis, G. & Mili, S. Translational regulation of protrusion-localized RNAs involves silencing and clustering after transport. Elife 8, doi:10.7554/eLife.44752 (2019).

29 Mili, S., Moissoglu, K. & Macara, I. G. Genome-wide screen reveals APC-associated RNAs enriched in cell protrusions. Nature 453, 115–119, doi:10.1038/nature06888 (2008).

30 Mardakheh, F. K. et al. Global Analysis of mRNA, Translation, and Protein Localization: Local Translation Is a Key Regulator of Cell Protrusions. Dev Cell 35, 344–357, doi:10.1016/j.devcel.2015.10.005 (2015).

31 Chrisafis, G. et al. Collective cancer cell invasion requires RNA accumulation at the invasive front. Proc Natl Acad Sci U S A 117, 27423–27434, doi:10.1073/pnas.2010872117 (2020).

32 Kuo, J. C., Han, X., Yates, J. R., 3rd & Waterman, C. M. Isolation of focal adhesion proteins for biochemical and proteomic analysis. Methods Mol Biol 757, 297–323, doi:10.1007/978-1-61779-166-6_19 (2012).

33 Kuang, R., Wang, Z., Xu, Q., Liu, S. & Zhang, W. Influence of mechanical stimulation on human dermal fibroblasts derived from different body sites. Int J Clin Exp Med 8, 7641–7647 (2015).

34 Byfield, F. J., Reen, R. K., Shentu, T. P., Levitan, I. & Gooch, K. J. Endothelial actin and cell stiffness is modulated by substrate stiffness in 2D and 3D. J Biomech 42, 1114–1119, doi:10.1016/j.jbiomech.2009.02.012 (2009).

35 Ma, W., Zheng, G., Xie, W. & Mayr, C. In vivo reconstitution finds multivalent RNA-RNA interactions as drivers of mesh-like condensates. Elife 10, doi:10.7554/eLife.64252 (2021).

36 Chen, K. H., Boettiger, A. N., Moffitt, J. R., Wang, S. & Zhuang, X. RNA imaging. Spatially resolved, highly multiplexed RNA profiling in single cells. Science 348, aaa6090, doi:10.1126/science.aaa6090 (2015).

37 om Dieck, S. et al. Direct visualization of newly synthesized target proteins in situ. Nature methods 12, 411–414, doi:10.1038/nmeth.3319 (2015).

38 Aratyn-Schaus, Y. & Gardel, M. L. Transient frictional slip between integrin and the ECM in focal adhesions under myosin II tension. Curr Biol 20, 1145–1153, doi:10.1016/j.cub.2010.05.049 (2010).

39 Dormond, O., Ponsonnet, L., Hasmim, M., Foletti, A. & Ruegg, C. Manganese-induced integrin affinity maturation promotes recruitment of alpha V beta 3 integrin to focal adhesions in endothelial cells: evidence for a role of phosphatidylinositol 3-kinase and Src. Thromb Haemost 92, 151–161, doi:10.1160/TH03-11-0728 (2004).

40 Moon, K. R. et al. Visualizing structure and transitions in high-dimensional biological data. Nat Biotechnol 37, 1482–1492, doi:10.1038/s41587-019-0336-3 (2019).

41 Kanchanawong, P. et al. Nanoscale architecture of integrin-based cell adhesions. Nature 468, 580–584, doi:10.1038/nature09621 (2010).

42 Panciera, T., Azzolin, L., Cordenonsi, M. & Piccolo, S. Mechanobiology of YAP and TAZ in physiology and disease. Nat Rev Mol Cell Biol 18, 758–770, doi:10.1038/nrm.2017.87 (2017).

43 Clark, K., Langeslag, M., Figdor, C. G. & van Leeuwen, F. N. Myosin II and mechanotransduction: a balancing act. Trends Cell Biol 17, 178–186, doi:10.1016/j.tcb.2007.02.002 (2007).

44 Kular, J. K., Basu, S. & Sharma, R. I. The extracellular matrix: Structure, composition, age-related differences, tools for analysis and applications for tissue engineering. J Tissue Eng 5, 2041731414557112, doi:10.1177/2041731414557112 (2014).

45 Jain, S. et al. ATPase-Modulated Stress Granules Contain a Diverse Proteome and Substructure. Cell 164, 487–498, doi:10.1016/j.cell.2015.12.038 (2016).

46 Guillen-Boixet, J. et al. RNA-Induced Conformational Switching and Clustering of G3BP Drive Stress Granule Assembly by Condensation. Cell 181, 346–361 e317, doi:10.1016/j.cell.2020.03.049 (2020).

47 Ivanov, P., Kedersha, N. & Anderson, P. Stress Granules and Processing Bodies in Translational Control. Cold Spring Harb Perspect Biol 11, doi:10.1101/cshperspect.a032813 (2019).

48 Hubstenberger, A. et al. P-Body Purification Reveals the Condensation of Repressed mRNA Regulons. Mol Cell 68, 144–157 e145, doi:10.1016/j.molcel.2017.09.003 (2017).

49 Youn, J. Y. et al. Properties of Stress Granule and P-Body Proteomes. Mol Cell 76, 286–294, doi:10.1016/j.molcel.2019.09.014 (2019).

50 Sabari, B. R., Dall’Agnese, A. & Young, R. A. Biomolecular Condensates in the Nucleus. Trends Biochem Sci 45, 961–977, doi:10.1016/j.tibs.2020.06.007 (2020).

51 Cho, W. K. et al. Mediator and RNA polymerase II clusters associate in transcription-dependent condensates. Science 361, 412–415, doi:10.1126/science.aar4199 (2018).

52 Fox, A. H., Nakagawa, S., Hirose, T. & Bond, C. S. Paraspeckles: Where Long Noncoding RNA Meets Phase Separation. Trends Biochem Sci 43, 124–135, doi:10.1016/j.tibs.2017.12.001 (2018).

53 Berkovits, B. D. & Mayr, C. Alternative 3’ UTRs act as scaffolds to regulate membrane protein localization. Nature 522, 363–367, doi:10.1038/nature14321 (2015).

54 Mateu-Regue, A. et al. Single mRNP Analysis Reveals that Small Cytoplasmic mRNP Granules Represent mRNA Singletons. Cell Rep 29, 736–748 e734, doi:10.1016/j.celrep.2019.09.018 (2019).

55 Yang, P. et al. G3BP1 Is a Tunable Switch that Triggers Phase Separation to Assemble Stress Granules. Cell 181, 325–345 e328, doi:10.1016/j.cell.2020.03.046 (2020).

56 Van Treeck, B. et al. RNA self-assembly contributes to stress granule formation and defining the stress granule transcriptome. Proc Natl Acad Sci U S A 115, 2734–2739, doi:10.1073/pnas.1800038115 (2018).

57 Roden, C. & Gladfelter, A. S. RNA contributions to the form and function of biomolecular condensates. Nat Rev Mol Cell Biol 22, 183–195, doi:10.1038/s41580-020-0264-6 (2021).

58 Chicurel, M. E., Singer, R. H., Meyer, C. J. & Ingber, D. E. Integrin binding and mechanical tension induce movement of mRNA and ribosomes to focal adhesions. Nature 392, 730–733, doi:10.1038/33719 (1998).

59 Wang, T., Hamilla, S., Cam, M., Aranda-Espinoza, H. & Mili, S. Extracellular matrix stiffness and cell contractility control RNA localization to promote cell migration. Nat Commun 8, 896, doi:10.1038/s41467-017-00884-y (2017).

60 Willett, M., Pollard, H. J., Vlasak, M. & Morley, S. J. Localization of ribosomes and translation initiation factors to talin/beta3-integrin-enriched adhesion complexes in spreading and migrating mammalian cells. Biol Cell 102, 265–276, doi:10.1042/BC20090141 (2010).

61 Huttelmaier, S. et al. Spatial regulation of beta-actin translation by Src-dependent phosphorylation of ZBP1. Nature 438, 512–515, doi:10.1038/nature04115 (2005).

62 Vejnar, C. E. & Giraldez, A. J. LabxDB: versatile databases for genomic sequencing and lab management. Bioinformatics 36, 4530–4531, doi:10.1093/bioinformatics/btaa557 (2020).

63 Aken, B. L. et al. Ensembl 2017. Nucleic Acids Res 45, D635–D642, doi:10.1093/nar/gkw1104 (2017).

64 Vejnar, C. E.

65 Jiang, H., Lei, R., Ding, S. W. & Zhu, S. Skewer: a fast and accurate adapter trimmer for next-generation sequencing paired-end reads. BMC Bioinformatics 15, 182, doi:10.1186/1471-2105-15-182 (2014).

66 Dobin, A. et al. STAR: ultrafast universal RNA-seq aligner. Bioinformatics 29, 15–21, doi:10.1093/bioinformatics/bts635 (2013).

67 Smit, A. F., Hubley, R. & Green, P. (1996).

68 Hofacker, I. (1994).

69 Lorenz, R. & Bernhart, S. Höner zu Siederdissen C et al (2011) ViennaRNA package 2.0. Algorithms Mol Biol 6, 26.

70 Moffitt, J. R. et al. High-throughput single-cell gene-expression profiling with multiplexed error-robust fluorescence in situ hybridization. Proc Natl Acad Sci U S A 113, 11046–11051, doi:10.1073/pnas.1612826113 (2016).

71 Rouillard, J. M., Zuker, M. & Gulari, E. OligoArray 2.0: design of oligonucleotide probes for DNA microarrays using a thermodynamic approach. Nucleic Acids Res 31, 3057–3062, doi:10.1093/nar/gkg426 (2003).

72 Camacho, C. et al. BLAST+: architecture and applications. BMC Bioinformatics 10, 421, doi:10.1186/1471-2105-10-421 (2009).

73 Horzum, U., Ozdil, B. & Pesen-Okvur, D. Step-by-step quantitative analysis of focal adhesions. MethodsX 1, 56–59, doi:10.1016/j.mex.2014.06.004 (2014).

74 Wang, S., Moffitt, J. R., Dempsey, G. T., Xie, X. S. & Zhuang, X. Characterization and development of photoactivatable fluorescent proteins for single-molecule-based superresolution imaging. Proc Natl Acad Sci U S A 111, 8452–8457, doi:10.1073/pnas.1406593111 (2014).

75 Liu, M. et al. Multiplexed imaging of nucleome architectures in single cells of mammalian tissue. Nat Commun 11, 2907, doi:10.1038/s41467-020-16732-5 (2020).

76 Stuart, T. et al. Comprehensive Integration of Single-Cell Data. Cell 177, 1888–1902 e1821, doi:10.1016/j.cell.2019.05.031 (2019).

77 Tom Dieck, S. et al. Direct visualization of newly synthesized target proteins in situ. Nat Methods 12, 411–414, doi:10.1038/nmeth.3319 (2015).

78 Kiema, T. et al. The molecular basis of filamin binding to integrins and competition with talin. Mol Cell 21, 337–347, doi:10.1016/j.molcel.2006.01.011 (2006).

79 Heuze, M. L. et al. ASB2 targets filamins A and B to proteasomal degradation. Blood 112, 5130–5140, doi:10.1182/blood-2007-12-128744 (2008).

80 Iwamoto, D. V. et al. Structural basis of the filamin A actin-binding domain interaction with F-actin. Nat Struct Mol Biol 25, 918–927, doi:10.1038/s41594-018-0128-3 (2018).

81 Kumar, A. et al. Filamin A mediates isotropic distribution of applied force across the actin network. J Cell Biol 218, 2481–2491, doi:10.1083/jcb.201901086 (2019).

